# Gaze stabilisation behaviour is anisotropic across visual field locations in zebrafish

**DOI:** 10.1101/754408

**Authors:** Florian A. Dehmelt, Rebecca Meier, Julian Hinz, Takeshi Yoshimatsu, Clara A. Simacek, Kun Wang, Tom Baden, Aristides B. Arrenberg

## Abstract

Many animals have large visual fields, and sensory circuits may sample those regions of visual space most relevant to behaviours such as gaze stabilisation and hunting. Despite this, relatively small displays are often used in vision neuroscience. To sample stimulus locations across most of the visual field, we built a spherical stimulus arena with 14,848 independently controllable LEDs, measured the optokinetic response gain of immobilised zebrafish larvae, and related behaviour to previously published retinal photoreceptor densities. We measured tuning to steradian stimulus size and spatial frequency, and show it to be independent of visual field position. However, zebrafish react most strongly and consistently to lateral, nearly equatorial stimuli, consistent with previously reported higher spatial densities in the central retina of red, green and blue photoreceptors. Upside-down experiments suggest further extra-retinal processing. Our results demonstrate that motion vision circuits in zebrafish are anisotropic, and preferentially monitor areas with putative behavioural relevance.

**Author summary:** The visual system of larval zebrafish mirrors many features present in the visual system of other vertebrates, including its ability to mediate optomotor and optokinetic behaviour. Although the presence of such behaviours and some of the underlying neural correlates have been firmly established, previous experiments did not consider the large visual field of zebrafish, which covers more than 160° for each eye. Given that different parts of the visual field likely carry unequal amount of behaviourally relevant information for the animal, this raises the question whether optic flow is integrated across the entire visual field or just parts of it, and how this shapes behaviour such as the optokinetic response. We constructed a spherical LED arena to present visual stimuli almost anywhere across their visual field, while tracking horizontal eye movements. By displaying moving gratings on this LED arena, we demonstrate that the optokinetic response, one of the most prominent visually induced behaviours of zebrafish, indeed strongly depends on stimulus location and stimulus size, as well as on other parameters such as the spatial and temporal frequency of the gratings. This location dependence is consistent with areas of high retinal photoreceptor densities, though evidence suggests further extraretinal processing.

## Introduction

The layout of the retina, and the visual system as a whole, evolved to serve specific behavioural tasks that animals perform to survive in their respective habitats. A well-known example is the position of the eyes in the head which varies between hunting animals (frontal eyes) and animals that frequently need to avoid predation (lateral eyes) (2). Hunting animals keep the prey within particular visual field regions to maximize behavioural performance (3–5). To avoid predation, however, it is useful to observe a large proportion of visual space, especially those regions in which predators are most likely to appear (6, 7). The ecological significance of visual stimuli thus depends on their location within the visual field, and it is paralleled by non-uniform processing channels across the retina. This non-uniformity manifests as an *area centralis* or a fovea in many species, which is a region of heightened photoreceptor density in the central retina and serves to increase visual performance in the corresponding visual field regions. Photoreceptor densities put a direct physical limit on performance parameters such as spatial resolution (8, 9). In addition to these restrictions mediated by the peripheral sensory circuitry, an animal’s use of certain visual field regions is also affected by behaviour-specific neural pathways, e.g. pathways dedicated to feeding and stabilisation behaviour. The retinal and extra-retinal circuit anisotropies can in turn effect a dependence of behavioural performance on visual field location (3, 4, 10–14).

Investigating behavioural performance limits and non-uniformities can offer insights into the processing capabilities and ecological adaptations of vertebrate brains, especially if they can be studied and quantitatively understood at each processing step. The larval zebrafish is a promising organism for such an endeavour, since its brain is small and a wide array of experimental techniques is available (15, 16). Zebrafish are lateral-eyed animals and have a large visual field, which covers 163° per eye (17). Their retina contains four different cone photoreceptor types (18), each distributed differently across the retina. UV photoreceptors are densest in the ventro-temporal retina (*area temporalis ventralis*), whereas the red, green and blue photoreceptors cover more central retinal regions (13).

Zebrafish larvae perform a wide range of visually mediated behaviours, ranging from prey capture (19) and escape behaviour (20) to stabilisation behaviour (21, 22), however the importance of stimulus location within the visual field for the execution of the respective behaviours has only recently been recognized and is still not well understood (3) (13, 23, 24).

During visually mediated stabilisation behaviours, such as optokinetic and optomotor responses, animals move their eyes and bodies, respectively, in order to stabilize the retinal image and/or the body position relative to the visual surround. The optokinetic response (OKR) consists of reflexively executed stereotypical eye movements, in which phases of stimulus “tracking” (slow phase) are interrupted by quick phases **(Figure 1a)**. In the quick phases, eye position is reset by a saccade in the direction opposite to stimulus motion. In humans, optokinetic responses are strongest in the central visual field (25). Furthermore, lower visual field locations of the stimulus evoke stronger OKR than upper visual field locations, which likely represents an adaptation to the rich optic flow information available from the structures on the ground in the natural environments of primates (10, 26).

**Figure 1.**
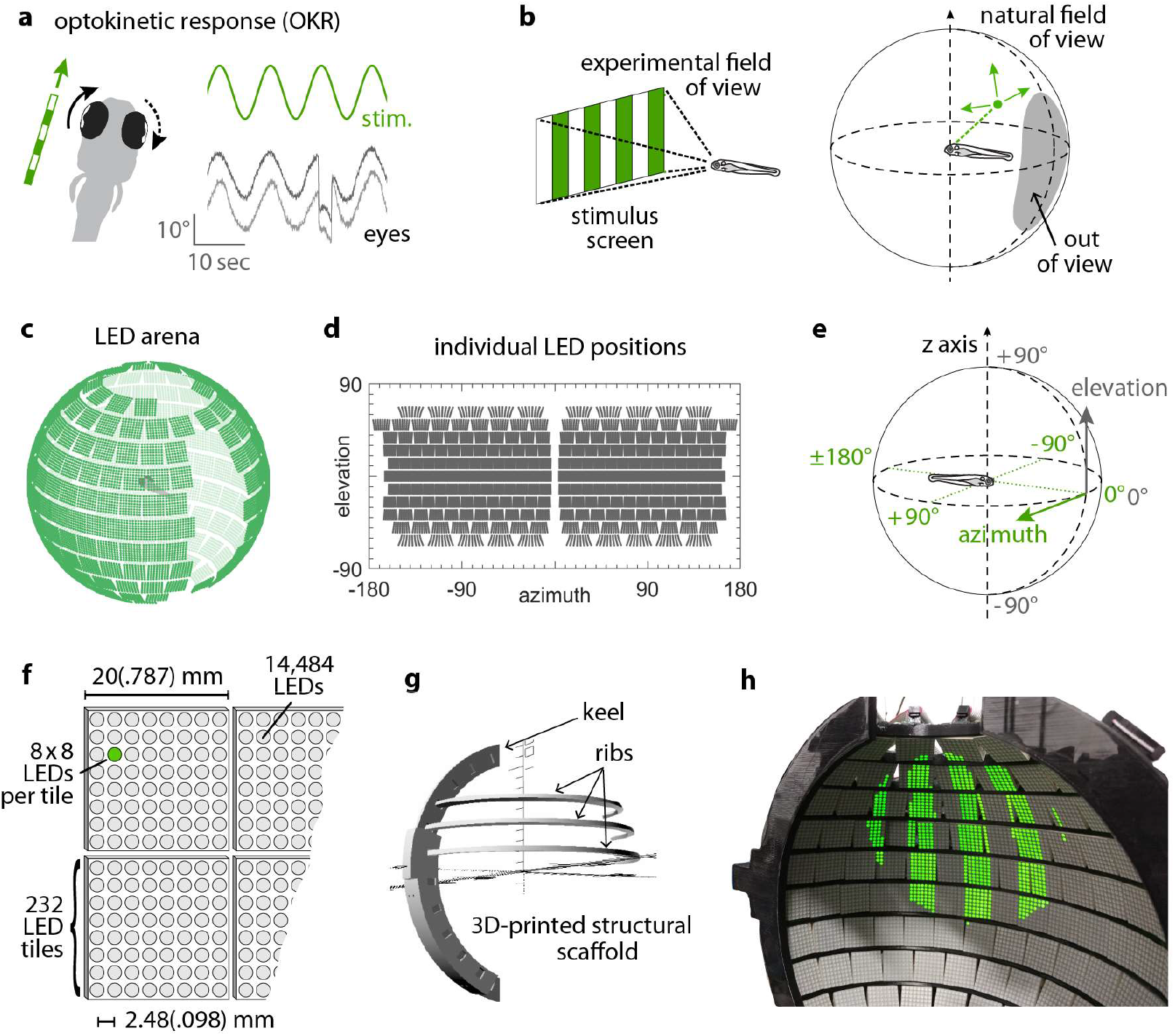
Presenting visual stimuli across the visual field. (a) When presented with a horizontal moving stimulus pattern, zebrafish larvae exhibit optokinetic response (OKR) behaviour, where eye movements track stimulus motion to minimise retinal slip. Its slow phase is interrupted by intermittent saccades, and even if only one eye is stimulated (solid arrow), the contralateral eye is indirectly yoked to move along (dashed arrow). (b) Often, experiments on visuomotor behaviour such as OKR sample only a small part of the visual field, whether horizontally or vertically. As different spatial directions may carry different behavioural importance, an ideal stimulation setup should cover all or most of the animal’s visual field. For zebrafish larvae, this visual field can be represented by an almost complete unit sphere. (c) We arranged 232 LED tiles with 64 LEDs each across a spherical arena, such that 14,484 LEDs (green dots) covered nearly the entire visual field. (d) The same individual positions, shown in geographic coordinates. Each circle represents a single LED. Each cohesive group of eight-by-eight circles corresponds to the 64 LEDs contained in a single tile. (e) To identify LED and stimulus locations, we use Up-East-North geographic coordinates: Azimuth *α* describes the horizontal angle, which is zero in front of the animal and, when seen from above, increases for rightward position. Elevation *β* refers to the vertical angle, which is zero throughout the plane containing the animal, and positive above. (f) The spherical arena is covered in flat square tiles carrying 64 green LEDs each. (g) Its structural backbone is made of a 3D-printed keel and ribs. Left and right hemispheres were constructed as separate units. (h) Across 85-90% of the visual field, we can then present horizontally moving bar patterns of different location, frequency and size to evoke OKR. **Figure 1–figure supplement 1.** A spherical LED arena to present visual stimuli across the visual field.

In zebrafish larvae, OKR behaviour has been used extensively to assess visual function in genetically altered animals (27–29). OKR tuning to the velocity, frequency, and contrast of grating stimuli has been measured (30–33), and, more recently, zebrafish are used as a model for investigating vertebrate sensorimotor transformations (21, 34). While zebrafish can distinguish rotational from translational optic flow to evoke appropriate optokinetic and optomotor responses (21, 35, 36), it is still unclear which regions of the visual field zebrafish preferentially observe in these behaviours. The aquatic lifestyle, in combination with the preferred swimming depths (37), might cause the lower visual field to contain less relevant information when compared to terrestrial animals. This in turn might result in behavioural biases to other – more informative – visual field regions. A corresponding systematic behavioural quantification in zebrafish or other aquatic species, which would relate OKR behaviour to naturally occurring motion statistics and the underlying neuronal representations in retina and retino-recipient brain structures, has been prevented by technical limitations. Specifically, little is known about (i) the dependence of OKR gain on stimulus location or (ii) on stimulus sizes, (iii) possible interactions between stimulus location, size and frequency, (iv) putative asymmetries between the left and right hemispheres of the visual field, and (v) the relationship between a putative dependence of OKR on stimulus location and zebrafish retinal architecture.

In other species with large visual fields, such as Drosophila, full-surround stimulation setups have been used successfully (1, 38, 39), but to date, none has been used for fish. This is at least partly due to their aquatic environment and the associated difficulties regarding the refraction of stimulus light at the air-water interface. Such distortions of shape can be partially compensated by pre-emptively altering the shape of the stimulus. However, using regular computer screens or video projection, the resulting luminance profiles remain anisotropic, potentially biasing the response toward brighter locations. Additionally, most stimulus arenas cannot easily be combined with the recording of neural activity, e.g., via calcium imaging, as stimulus light and calcium fluorescence overlap in both the spectral and time domains. These challenges must be overcome to enable full-field visual stimulation in zebrafish neurophysiology experiments **(Figure 1b)**.

Here, we present a novel visual stimulus arena for aquatic animals, which covers almost the entire surround of the animal, and use it to characterize the anisotropy of the zebrafish OKR across different visual field locations as well as the tuning to stimulus size, spatial frequency and leftside versus rightside stimulus locations. We find that the OKR is mostly symmetric across both eyes and driven most strongly by lateral stimulus locations. These stimulus locations approximately correspond to a retinal region of increased photoreceptor density. By rotating the experimental setup and/or the animal, our control experiments revealed that additional extra-retinal determinants of OKR drive exist as well. Our characterization of OKR drive across the visual field will help inform bottom-up models of the vertebrate neural pathways underlying optokinetic and other visual behaviour.

## Results

### Spherical LED arena allows presentation of stimuli across the visual field

By combining 3D printing with electronic solutions developed in *Drosophila* vision research, we constructed a spherical stimulus arena containing 14,848 individual LEDs covering over 90% of the visual field of zebrafish larvae **(Figure 1c, Supplementary File 1)**. Using infrared illumination via an optical pathway coupled into the sphere **(Figure 1– figure supplement 1a-b)**, we tracked eye movements of larval zebrafish during presentation of visual stimuli (40).

To avoid stimulus aberrations at the air-to-water interface, we designed a nearly spherical glass bulb containing fish and medium **(Figure 1–figure supplement 1c-d)**. With this design, stimulus light from the surrounding arena is virtually not refracted (light is orthogonal to the air-to-water interface), and reaches the eyes of the zebrafish larva in a straight line. Thus, no geometric corrections are required during stimulus design **(Supplementary Code 1)**, and stimulus luminance is expected to be nearly isotropic across the visual field. We additionally designed the setup to minimise visual obstruction, and developed a new embedding technique to immobilise the larva at the tip of a narrow glass triangle (see **Methods**). In almost all possible positions, fish can thus perceive stimuli without interference. The distance between most of the adjacent LED pairs is smaller than the photoreceptor spacing in the larval retina (8, 41), resulting in a good spatial resolution across the majority of the spherical arena surface (see detailed discussion in **Supplementary File 2**). As flat square LED tiles cannot be perfectly arranged on a spherical surface, small triangular gaps are unavoidable. More importantly, several gaps in LED coverage, resulting from structural elements of the arena, were restricted mainly to the back, the top, and bottom of the animal. The “keel” behind and in front of the fish supports the horizontal “ribs”, and the circular openings in the top and bottom accommodate the optical path for eye tracking or scanning microscopy **(Supplementary File 3)**.

### Stimulus position dependence of the optokinetic response

Horizontally moving vertical bars reliably elicit OKR in zebrafish larvae (42). We used a stimulus which rotated clock- and counter-clockwise with a sinusoidal velocity pattern (velocity amplitude 12.5 degree/sec, frequency of the velocity envelope 0.1 Hz, spatial frequency 0.06 cycles/degree, **Figure 2a**). OKR performance was calculated by measuring the amplitude of the resulting OKR slow-phase eye movements after the saccades had been removed **(Figure 2b, Figure 2–figure supplement 1, Supplementary Code 3, Methods)**. The OKR gain then corresponds to the speed of the slow-phase eye movements divided by the speed of the stimulus (which is equivalent to the ratio of the eye position and stimulus position amplitudes). In addition to traditional full-field stimulation, our arena can display much smaller stimuli in different parts of the visual field. These novel stimuli evoked reliable OKR even at remote stimulus locations **(Figure 2c, Figure 2– figure supplement 2)**, and thus allowed us to investigate previously inaccessible behavioural parameters. We excluded any trials from data analysis that showed other behaviours (e.g. drifts and ongoing spontaneous eye movements) superimposed on OKR **(Figure 2–figure supplement 2)**. In addition to the characteristic eye traces **(Figure 2c)**, Bode plots of the magnitude and phase shift relative to the stimulus qualitatively match previously reported zebrafish response to full-field stimulation **(Figure 5–figure supplement 1**, (42)), further confirming that the behaviour we observe is indeed OKR.

**Figure 2.**
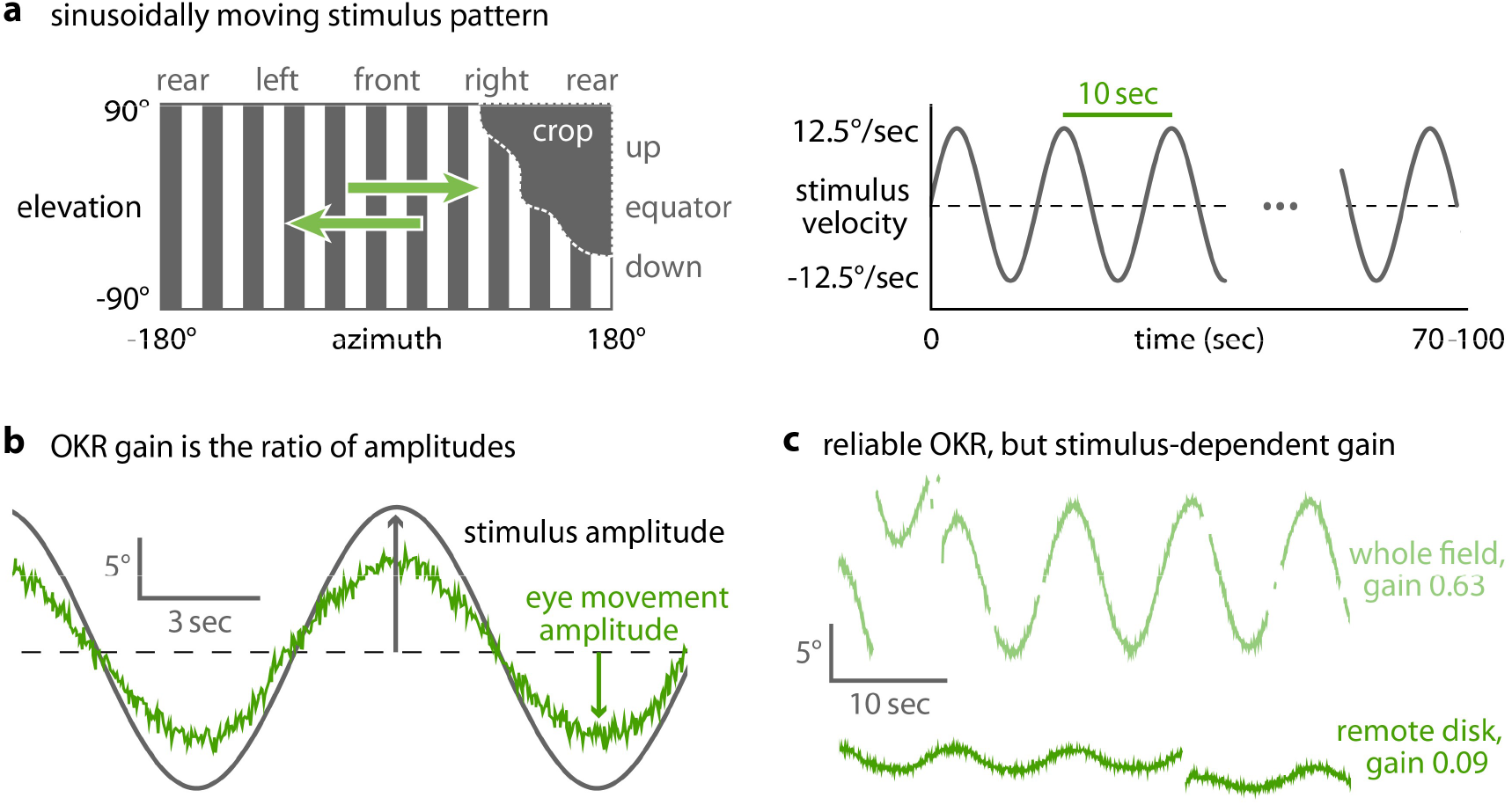
OKR gain is inferred from a piece-wise fit to the slow phase of tracked eye movements. (a) We present a single pattern of horizontally moving bars to evoke OKR, and crop it by superimposing a permanently dark area of arbitrary size and shape (left). Its velocity follows a sinusoidal time course, repeating every 10 seconds for a total of 100 seconds for each stimulus phase (right). (b) OKR gain is the amplitude of eye movement (grey trace) relative to the amplitude of the sinusoidal stimulus (green trace). The OKR gain is often well below 1, e.g. for high stimulus velocities as used here (up to 12.5°/s). (c) Even small stimuli are sufficient to elicit reliable OKR, though gains are low if stimuli appear in a disfavoured part of the visual field. Shown here are responses to a whole-field stimulus (top) and to a disk-shaped stimulus in the extreme upper rear (bottom, **Figure 2–figure supplement 2**). **Figure 2–figure supplement 1.** Saccades are cropped before OKR gain is measured. **Figure 2–figure supplement 2.** Even at remote stimulus locations, fish exhibit reliable OKR behaviour; trials with mixed behaviour are excluded.

To quantify position tuning, we cropped the presented gratings **(Figure 2a)** to a disk-shaped area of constant size, centred on one of 38 nearly equidistant parts of the visual field **(Figure 3a, Table 1, Video 1, Video 2)**. The distribution of positions was symmetric between the left and right, upper and lower, as well as front and rear hemispheres, with some stimuli falling right on the edge between two hemispheres. As permanent asymmetries in a stimulus arena or in its surroundings could affect OKR gain, we repeated our experiments in a second group of larvae after rotating the arena by 180 degrees **(Figure 4a-b)**, then matched the resulting pairs of OKR gains during data analysis (see **Methods, Figure 4e-f**). Any remaining asymmetries in the OKR distributions should result from biological lateralisation.

**Figure 3.**
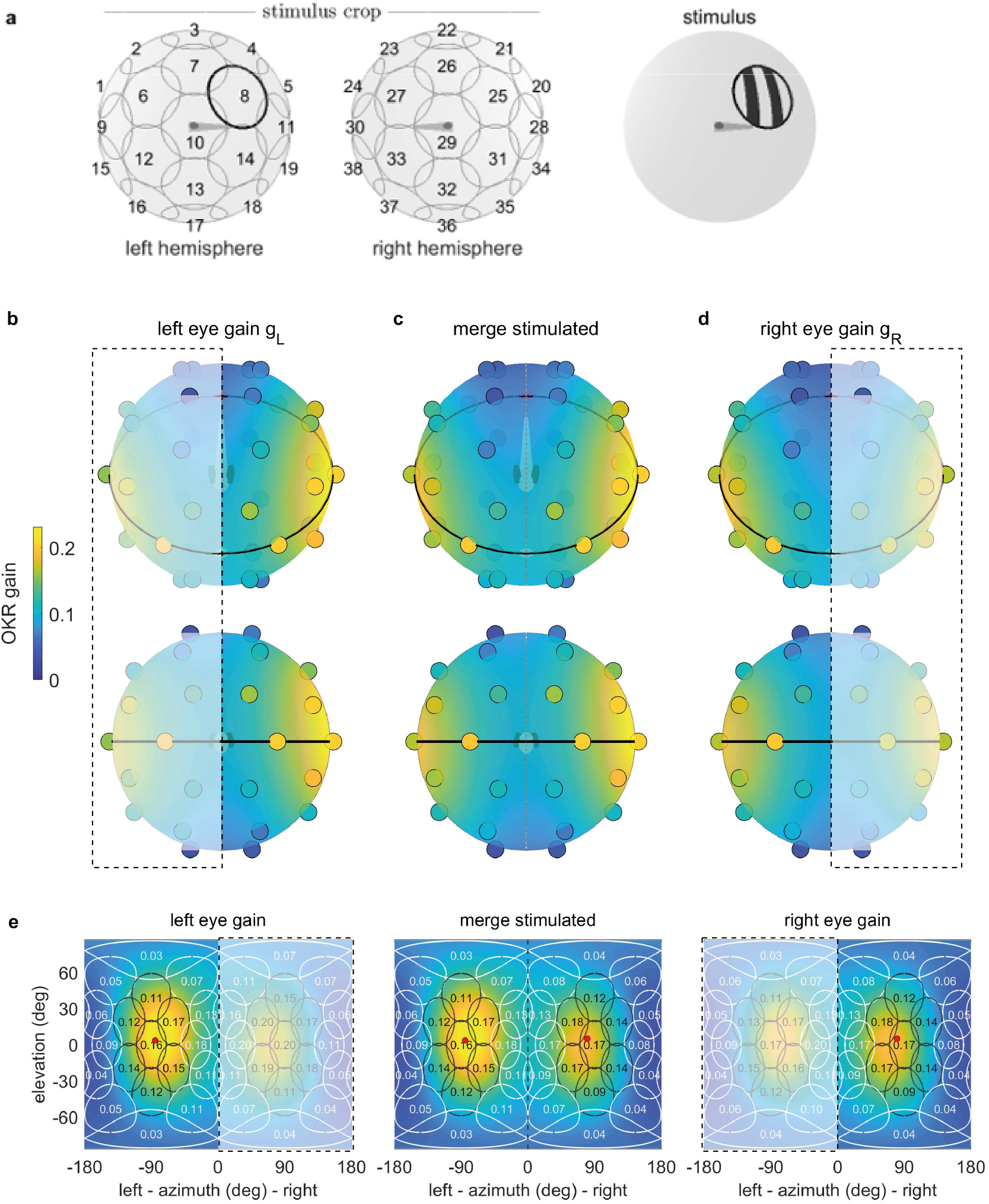
OKR gain depends on stimulus location. (a) The stimulus is cropped to a disk-shaped area 40 degrees in diameter, centred on one of 38 nearly equidistant locations (cf. Table 1) across the entire visual field (left), to yield 38 individual stimuli (right). (b-d) Dots reveal the location of stimulus centres D1-D38. Their colour indicates the average OKR gain across individuals and trials, corrected for external asymmetries. Surface colour of the sphere displays the best von-Mises Fisher fit to the discretely sampled OKR data. Top row: OKR gain of the left eye (b), right eye (d), and the merged data including only direct stimulation of either eye (c), shown from an oblique, rostrodorsal angle. Bottom row: same, but shown directly from the front. OKR gain is significantly higher for lateral stimulus locations and lower across the rest of the visual field. The spatial distribution of OKR gains is well explained by the bimodal sum of two von-Mises Fisher distributions. (e) Mercator projections of OKR gain data shown in panels (b-d). White and grey outlines indicate the area covered by each stimulus type. Numbers indicate average gain values for stimuli centred on this location. Red dots show mean eye position during stimulation. Dashed outline and white shading on panels (b, d, e) indicate indirect stimulation via yoking, i.e., stimuli not directly visible to either the left or right eye. Data from n=7 fish for the original configuration and n=5 fish for the rotated arena. **Figure 3–figure supplement 1.** Gaps in the arena do not bias OKR behaviour. **Figure 3–figure supplement 2.** OKR beating field and average eye position are independent of stimulus location. **Figure 3–figure supplement 3.** Vertical eye position under upright and upside-down embedding. **Figure 3–figure supplement 4.** Yoking indices are biased by reflections within the arena. **Figure 3–figure supplement 5.** Reflections alter perceived yoking indices across arena types.

**Figure 4.**
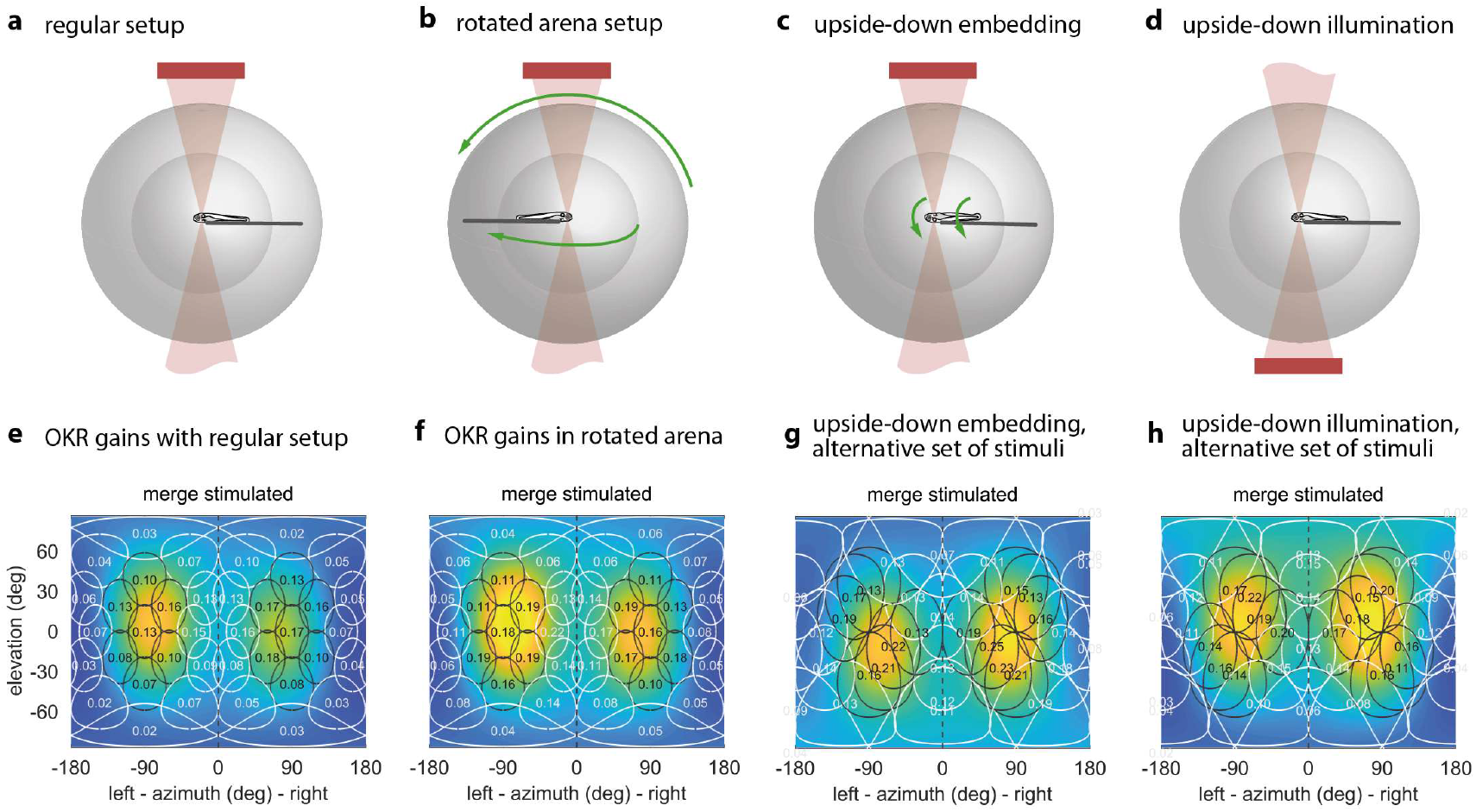
The OKR is biased towards upper environmental elevations irrespective of fish orientation. (a) Regular arena setup. (b) Arena can be tilted 180 degrees so front and rear, upper and lower LED positions are swapped. The bulb holder moves accordingly, so from the perspective of the fish, left and right, upper and lower LEDs are swapped. (c) Upside-down embedding setup. (d) Setup with inverted optical path, including illumination. (e-h) Results in body-centred coordinates, where positive elevations refer to dorsal positions, for the four setups shown in (a-d). (g,h) Experiments with presentation of a less regularly distributed set of stimuli, cropped to disks of 64 degrees polar angle instead of the 40 degrees used in (a,b,e,f). (g) Fish embedded upside-down exhibit a slight preference for stimuli below the body-centred equator, i.e., positions slightly ventral to their body axis. (h) Fish embedded upright, as in (a). To account for environmental asymmetries such as arena anisotropies, we combined the data underlying (e) and (f) to obtain **Figure 3b-e** (see **Methods**). Data from (e) n=7, (f) n=5, (g) n=3, (h) n=10 fish. **Figure 4–figure supplement 1.** Individual fish exhibit weak and broadly-distributed biases towards the left or right half of their visual field.

**Table 1.** Stimulus parameters (position dependence). These stimuli consisted of a horizontally moving grating, cropped with a disk-shaped stimulus mask, and presented in one of 38 different locations across the visual field. Parameters as in **Table 1**. Results shown in **Figure 3** and **Figure 4e-f**.

| type | $\alpha$ | $\beta$ | $\delta$ | SF | TF | v | T |
| --- | --- | --- | --- | --- | --- | --- | --- |
| D1 | -16 | 25 | 64 | 0.0611 | 0.7639 | 12.5 | 10 |
| D2 | -35 | 53 | 64 | 0.0611 | 0.7639 | 12.5 | 10 |
| D3 | -90 | 74 | 64 | 0.0611 | 0.7639 | 12.5 | 10 |
| D4 | -145 | 53 | 64 | 0.0611 | 0.7639 | 12.5 | 10 |
| D5 | -164 | 25 | 64 | 0.0611 | 0.7639 | 12.5 | 10 |
| D6 | -60 | 19 | 64 | 0.0611 | 0.7639 | 12.5 | 10 |
| D7 | -90 | 39 | 64 | 0.0611 | 0.7639 | 12.5 | 10 |
| D8 | -120 | 19 | 64 | 0.0611 | 0.7639 | 12.5 | 10 |
| D9 | -30 | 0 | 64 | 0.0611 | 0.7639 | 12.5 | 10 |
| D10 | -90 | 0 | 64 | 0.0611 | 0.7639 | 12.5 | 10 |
| D11 | -150 | 0 | 64 | 0.0611 | 0.7639 | 12.5 | 10 |
| D12 | -60 | -19 | 64 | 0.0611 | 0.7639 | 12.5 | 10 |
| D13 | -90 | -39 | 64 | 0.0611 | 0.7639 | 12.5 | 10 |
| D14 | -120 | -19 | 64 | 0.0611 | 0.7639 | 12.5 | 10 |
| D15 | -16 | -25 | 64 | 0.0611 | 0.7639 | 12.5 | 10 |
| D16 | -35 | -53 | 64 | 0.0611 | 0.7639 | 12.5 | 10 |
| D17 | -90 | -74 | 64 | 0.0611 | 0.7639 | 12.5 | 10 |
| D18 | -145 | -53 | 64 | 0.0611 | 0.7639 | 12.5 | 10 |
| D19 | -164 | -25 | 64 | 0.0611 | 0.7639 | 12.5 | 10 |
| D20-D38 | same as D1-D19, but with positive azimuth $\alpha$ (right hemisphere of arena) | | | | | | |

To overcome our spatially discrete sampling, we then fit our data with a bimodal function comprised of two Gaussian-like two-dimensional distributions on the stimulus sphere surface (see **Methods, Supplementary Code 4**), to determine the location of highest OKR gain evoked by ipsilateral stimuli and contralateral stimuli, respectively. We observed significantly higher OKR gains in response to nearly lateral stimuli, and lower gains across the rest of the visual field **(Figure 3b-e)**. OKR was strongest for stimuli near an azimuth of 80.3 degrees and an elevation of 6.1 degrees for the left side (in body-centred coordinates), as well as −77.0 and −2.0 degrees for the right side – slightly rostral of the lateral meridian, and very close to the equator. Note that due to the fast stimulus speeds, the absolute slow phase eye velocities were high, while the OKR gain was relatively low. We chose such high stimulus speeds in order to minimize the experimental recording time needed to obtain reliable OKR measurements for each visual field location.

As our stimulus arena is not completely covered by LEDs **(Figure 1c, Figure 1d)**, some areas remain permanently dark. These could interfere with the perception of stimuli presented on adjacent LEDs. This is especially relevant as LED coverage is almost perfect for some stimulus positions (near the equator), whereas the size of triangular holes increases at others (towards the poles). We thus performed control experiments comparing the OKR gain evoked by a stimulus in a densely-covered part of the arena to the OKR gain evoked by same stimulus, but in the presence of additional dark triangular patches **(Figure 3–figure supplement 1a)**. We found no significant difference in OKR gain (**Figure 3–figure supplement 1c**, t-test, p<0.05). Additionally, we performed another series of control experiments using a dark shape mimicking the dark structural elements, the front “keel” of the arena **(Figure 3–figure supplement 1b)**. Again, we found no difference in OKR gain (**Figure 3–figure supplement 1c**, t-test, p<0.05), and thus ruled out that position dependence data was corrupted by incomplete LED coverage. Since the eyes were moving freely in our experiments, the range of eye positions during OKR, or so-called beating field (43), could have changed with stimulus position. We found that animals instead maintained similar median horizontal eye positions (e.g., left eye: −83.7±1.8 degrees, right eye: 80.3±1.9 degrees, average median ± standard deviation of medians, n=7 fish, **Figure 3–figure supplement 2**) even for the most peripheral stimulus positions.

A priori, it is unclear whether the sampling preference originates from the peculiarities of the sensory periphery in the eye, or the behavioural relevance inferred by central brain processing. The former would prioritise stimulus preference based on its position relative to the eye and, by extension, its representation on specific parts of the retina. The latter would prioritise stimulus preference based on its position relative to the environment, such as a predator approaching from the water surface. To distinguish both possible effects in the context of OKR, as well as to reveal any stimulus asymmetries accidentally introduced during the experiment, we performed control experiments with larvae embedded upside-down (i.e., with their dorsum towards the lower pole of the arena, **Figure 4c**). As a result of this upside-down embedding, world-centred and fish-centred coordinate systems were no longer identical, in that “up” in one is “down” in the other, and “left” becomes “right”. To facilitate comparisons across embedding types, all positions from here onwards are given relative to the visual field, and thus in fish-centred coordinates. Unexpectedly, the elevation of highest OKR gains relative to the eye changed from slightly above to slightly below the equator of the visual field when comparing upright to inverted fish **(Figure 4e, Figure 4g)**: When upright, azimuths and body-centred elevations of the peaks of the best fit to data were −67.8° and 8.4° for the left eye, as well as 73.1° and 6.2° for the right eye. When inverted, −88.8° and −1.2° for the left eye, as well as 80.0° and −12.2° for the right eye. These numbers were obtained from the gains of those eyes to which any given stimulus was directly visible. The results from **Figure 4e-f** were combined to correct **Figure 3** for external asymmetries (**Methods**); this is why the best-fit position reported for **Figure 4e** alone differs slightly from that reported above for **Figure 3b-e**. Because the set of visual stimuli presented to inverted fish stemmed from an earlier stimulus protocol with less even sampling of the visual field, a slight scaling of azimuths and elevations is expected. The consistent sign-change of the elevation, however, is not. We performed a permutation test in which embedding-direction labels were randomly swapped while stimulus-location labels were maintained, and the Gaussian-type fit to data was then repeated on each permuted dataset. This test confirmed that fish preferred upward (in environmental conditions) rather than dorsalward elevations (p < 0.05, **Supplementary Code 5**).

Adjustment by the fish of its vertical resting eye position between the upright and inverted body positions would have been a simple potential explanation for this result. However, time-lapse frontal microscopy images **(Methods)** ruled this out, since for both upside-up and upside-down embedding the eyes were inclined by an average of about 4 degrees towards the dorsum (3.5±1.0° for the left eye, 4.9±0.8° for the right eye, mean ± s.e.m., **Figure 3–figure supplement 3**). We also tested the influence of camera and infrared light (840 nm) positions **(Figure 4d)** – which in either case should have been invisible to the fish (44) – and found that they could indeed not explain the observed differences **(Figure 4h)**. In summary, the body-centred preferred location only flipped from slightly dorsal to slightly ventral in upside-down embedded fish **(Figure 4g)**, and thus remained virtually unchanged in environmental coordinates across all control experiments. Optokinetic stimulus location preference appears to be related to the behavioural relevance of these stimulus positions, and cannot merely be caused by retinal feedforward circuitry.

### Yoking of the non-stimulated eye

Almost all stimuli were presented monocularly – that is, in a position visible to only one of the two laterally located eyes. Without exception, zebrafish larvae responded with yoked movements of both the stimulated and unstimulated eye. To rule out reflections of stimuli within the arena, we performed a series of experiments in which the unstimulated side of the glass bulb had been covered with a matte, black sheet of plastic. Reflections on the glass-air interface would otherwise cause monocular stimuli (that should only be visible to the ipsilateral eye) to also be seen by the contralateral eye. Yoking indices (YI) were significantly different between the regular monocular setup (YI≈0.2) and the control setup (YI≈0.7) containing the black surface on the side of the unstimulated eye, confirming that yoking indices had been affected by reflections (**Figure 3–figure supplement 4**, an index of 1 indicating completely monocular eye movements, an index of 0 perfectly conjugate eye movements/yoking). This suggests that previously reported yoking partially depends on reflections of the stimulus pattern at the glass-to-air or water-to-air interface in our spherical setup and other commonly used stimulus arenas. We performed additional control experiments using a previously described setup (40) with four flat LCD screens for stimulus presentation in a different room. In these experiments, stimuli were presented monocularly or binocularly, and the unstimulated eye was either (i) stimulated with a stationary grating **(Figure 3–figure supplement 5a-b)**, (ii) shielded with a blank, white shield placed directly in front of the displays **(Figure 3–figure supplement 5c-d)** or (iii) shielded with a matte, black sheet of aluminium foil placed inside the petri dish (control for possible reflections on the Petri dish wall) **(Figure 3–figure supplement 5e-f)**. This experiment showed that yoking was much reduced (YI≈0.3) if the non-stimulated eye saw a stationary grating (i) instead of the white or black shields (ii-iii, YI≈0.1) or a binocular motion stimulus (YI≈0) (**Figure 3–figure supplement 5g-h**, p<0.05).

### Spatial asymmetries

As multiple previous studies reported left-right asymmetries in zebrafish visuomotor processing and behaviour other than OKR (45–47), we computed an asymmetry index *B* **(Methods)** to reveal whether zebrafish OKR is lateralised in individuals or across the population. We did not observe a general asymmetry between the response of the left and right eyes. Rather, our data is consistent with three distinct sources of asymmetry: individual bias towards one eye, shared bias across individuals, and asymmetries induced by the environment (including the experimental setup and stimulus arena). Through multivariate linear regression, we fit a linear model of asymmetries to our data **(Methods)**, which combined data from fish embedded upside-up **(Figure 4e)**, upside-down **(Figure 4g)** and data obtained with the arena rotated relative to the fish **(Figure 4f)**. Regression coefficients for external causes of asymmetry were similar to or smaller than those for biological causes **(Figure 4–figure supplement 1)**, and individual biases from fish to fish were broadly and symmetrically distributed from left to right (mean coefficient 3.7 ∙ 10^−4^ ± 120.0 ∙ 10^−4^ st. dev., n =15), so that no evidence was found for a strong and consistent lateralisation of OKR behaviour across animals.

Our results show that the OKR behaviour is mostly symmetric across both eyes, with individual fish oftentimes having a dominant eye due to seemingly random bias for one eye (lateralisation) across fish. Some of the observed asymmetries are consistent with external factors. Therefore, the OKR gains presented in **Figure 3** have been corrected in order to present only biologically meaningful differences **(Methods)**.

### Spatial frequency tuning of the optokinetic response is similar across visual field locations

We investigated the spatial frequency tuning of OKR behaviour across visual field positions by presenting 7 different spatial frequencies of the basic stimulus, each cropped into a planar angle of 40 degrees, at different visual field locations **(Figure 5a)**. Because we held the temporal frequency constant, stimulus velocity decreased whenever spatial frequency increased. These 7 disk-shaped stimuli were presented while centred on one of 6 possible locations in different parts of the visual field **(Figure 5c)**, with 3 locations on each hemisphere: one near the location of highest OKR gain as determined in our experiments on position dependence, one in a nasal location, and one in a lower temporal location. In total, we thus presented 42 distinct types of stimuli **(Table 2, Video 3)**. For each stimulus location and eye, the highest OKR gain was observed at a spatial frequency of 0.03 to 0.05 cycles/degree **(Figure 5d)**. We did not observe any strong modulation of frequency dependence by stimulus location.

**Figure 5.**
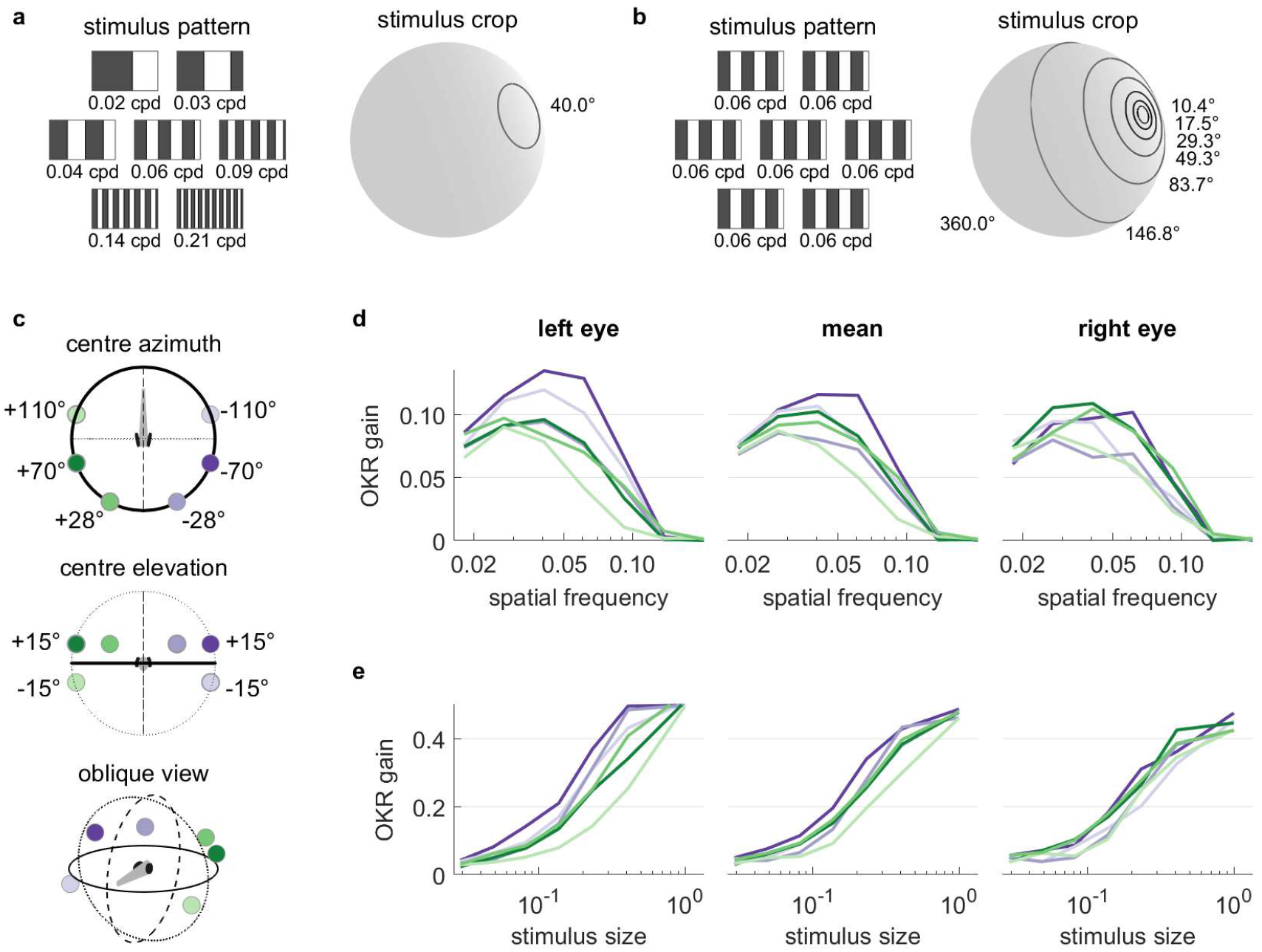
OKR tuning to spatial frequency is similar across different visual field locations. (a) Patterns with 7 different frequencies were cropped to disks of a single size. These disks were placed in 6 different locations for a total of 42 stimuli. cpd: cycles per degree. (b) Patterns with identical spatial frequencies were cropped to disks of 7 different sizes. These disks were also placed in 6 different locations for another set of 42 stimuli. Degrees indicate planar angles subtended by the stimulus outline, so 360° correspond to whole-field stimulation. (a, b) Displaying the entire actual pattern at the size of this figure would make the individual bars hard to distinguish. We thus only show a zoomed-in version of the patterns in which 45 out of 360 degrees azimuth are shown. (c) Coloured dots indicate the 6 locations on which stimuli from a and b were centred, shown from above (top), from front (middle), and from an oblique angle (bottom). (d) OKR gain is unimodally tuned to a wide range of spatial frequency (measured in cycles per degree). (e) OKR gain increases sigmoidally as the area covered by the visual stimulus increases logarithmically (a stimulus size of 1 corresponds to 100% of the spherical surface). (d-e) Colours correspond to the location of stimulus centres shown in (c). There is no consistent dependence on stimulus location of either frequency tuning or size tuning. Data from n=7 fish for frequency dependence and another n=7 fish for size dependence. **Figure 5–figure supplement 1.** Magnitude and phase shift of eye movements at different frequencies resemble those previously observed for zebrafish OKR.

**Table 2.** Stimulus parameters (frequency dependence). These stimuli consisted of a horizontally moving grating, cropped with a disk-shaped stimulus mask. At each location, 7 different spatial frequencies and thus velocities were used, while temporal frequency was held constant. Parameters and units as in Table 1. Results shown in **Figure 5**.

| type | $\alpha$ | $\beta$ | $\delta$ | SF | TF | v | T |
| --- | --- | --- | --- | --- | --- | --- | --- |
| **F1** | -70 | 15 | 40 | 0.0181 | 0.7639 | 42.188 | 10 |
| **F2** | -70 | 15 | 40 | 0.0272 | 0.7639 | 28.125 | 10 |
| **F3** | -70 | 15 | 40 | 0.0407 | 0.7639 | 18.750 | 10 |
| **F4** | -70 | 15 | 40 | 0.0611 | 0.7639 | 12.500 | 10 |
| **F5** | -70 | 15 | 40 | 0.0917 | 0.7639 | 8.3333 | 10 |
| **F6** | -70 | 15 | 40 | 0.1375 | 0.7639 | 5.5556 | 10 |
| **F7** | -70 | 15 | 40 | 0.2063 | 0.7639 | 3.7037 | 10 |
| **F8-F14** | same as V1-V7, but with azimuth $\alpha = -110$ and elevation $\beta = -15$ | | | | | | |
| **F15-F21** | same as V1-V7, but with azimuth $\alpha = -28$ and elevation $\beta = 15$ | | | | | | |
| **F22-F42** | same as A1-A21, but with positive azimuth $\alpha$ (right hemisphere) | | | | | | |

### Size dependence of the optokinetic response

It is unclear to what extent small stimuli are effective in driving OKR. We therefore employed a stimulus protocol with 7 OKR stimuli presented in differently sized areas on the sphere **(Figure 5b)**. Spatial and temporal frequencies were not altered, so bars appeared with the same width and velocity profile in all cases. These 7 disk-shaped stimuli were presented while centred on one of 6 possible locations, identical to those used to study frequency dependence **(Figure 5c)**, again yielding 42 unique stimuli **(Table 3, Video 4)**. Stimulus area size was chosen at logarithmic intervals, ranging from stimuli almost as small as the spatial resolution of the zebrafish retina, to stimuli covering the entire arena. Throughout this paper, the term “stimulus size” refers to the fractional area of the sphere surrounding the fish, in which the moving grating stimuli was presented. E.g. for a solid angle of 180°, the stimulus size is 50% and covers half of the surrounding space **(Fig. 5b)**. In line with many other psychophysical processes, OKR gain increased sigmoidally with the logarithm of stimulus size **(Figure 5e)**. Weak OKR behaviour was already observable in response to very small stimuli of 0.8 % (solid angle or “stimulus diameter” of 10.4°), and reached half-maximum performance at a stimulus size of roughly 25 % (solid angle: 120°). As was the case for spatial frequency dependence, we did not observe any strong modulation of size dependence by stimulus location, although OKR gains of the left eye appeared more dependent on stimulus location than those of the right eye.

**Table 3.** Stimulus parameters (size dependence). These stimuli consisted of a horizontally moving grating, cropped with a disk-shaped stimulus mask. At each location, disks with 7 different, logarithmically spaced areas were shown. Parameters and units as in **Table 1**. Results shown in **Figure 5**.

| type | $\alpha$ | $\beta$ | $\delta$ | SF | TF | v | T |
| --- | --- | --- | --- | --- | --- | --- | --- |
| **A1** | -70 | 15 | 3.58 | 0.0611 | 0.7639 | 12.5 | 10 |
| **A2** | -70 | 15 | 7.16 | 0.0611 | 0.7639 | 12.5 | 10 |
| **A3** | -70 | 15 | 14.3 | 0.0611 | 0.7639 | 12.5 | 10 |
| **A4** | -70 | 15 | 28.7 | 0.0611 | 0.7639 | 12.5 | 10 |
| **A5** | -70 | 15 | 57.9 | 0.0611 | 0.7639 | 12.5 | 10 |
| **A6** | -70 | 15 | 120 | 0.0611 | 0.7639 | 12.5 | 10 |
| **A7** | -70 | 15 | 360 | 0.0611 | 0.7639 | 12.5 | 10 |
| **A8-A14** | same as A1-A7, but with azimuth $\alpha = -110$ and elevation $\beta = -15$ | | | | | | |
| **A15-A21** | same as A1-A7, but with azimuth $\alpha = -28$ and elevation $\beta = 15$ | | | | | | |
| **A22-A42** | same as A1-A21, but with positive azimuth $\alpha$ (right hemisphere) | | | | | | |

### Optokinetic response gain covaries with retinal density of long-wave sensitive photoreceptors

We hypothesized that the non-uniform distribution of the OKR gain is related to the surface density of photoreceptors and investigated this using data from a recent study (13) on photoreceptor densities in explanted eye cups of 7-8 day old zebrafish larvae. As shown in **Figure 6b**, ultraviolet receptor density exhibits a clear peak in the upper frontal part of the visual field, whereas red, green and blue receptors **(Figure 6a)** are most concentrated across a wider region near the intersection of the equator and lateral meridian, with a bias to the upper visual field (in body coordinates). For comparison, density maps in retinal coordinates, not body coordinates, are shown in **Figure 6–figure supplement 1**, and **Figure 6c** shows total photoreceptor density across all types. To register our OKR gain data onto the photoreceptor density maps, we took the average eye position into account, which was located horizontally at −84.8±6.2 degrees azimuth for the left and 80.1±6.5 deg for the right eye (mean±st.dev., n=7 fish), and vertically at 3.5±3.2 degrees elevation for the left and 4.9±2.7 deg for the right eye (n=10 fish). For green, blue and especially red receptors, the visual space location in which OKR gain is maximal (cf, **Figure 3e**) coincides with a retinal region of near-maximum photoreceptor density (white ring in **Figure 6**). For ultraviolet receptors **(Figure 6b)**, there is no strong correlation between photoreceptor density and OKR gain.

**Figure 6.**
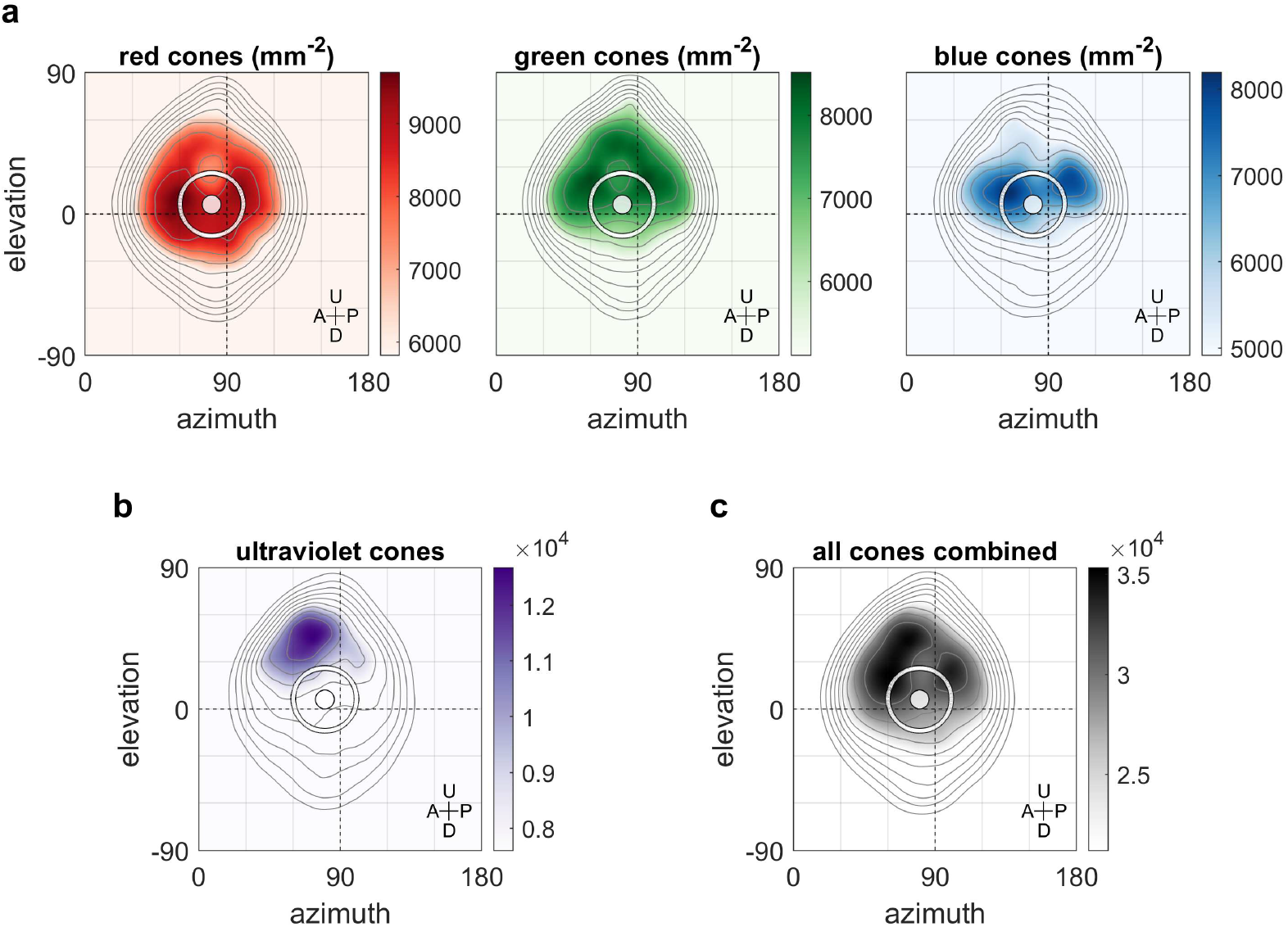
Maximum OKR gain is consistent with high photoreceptor densities in the retina. Contour lines show retinal photoreceptor density determined by optical measurements of explanted eye cups of 7-8 dpf zebrafish larvae, at increments of 10% of maximum density. Data shown in visual space coordinates relative to the body axis, i.e., 90° azimuth and 0° elevation corresponds to a perfectly lateral direction. To highlight densely covered regions, densities from half-maximum to maximum are additionally shown in shades of colour. Solid white circles indicate the location of maximum OKR gain inferred from experiments of type D in 5-7dpf larvae **(Figure 3)**. White outlines indicate the area that would be covered by a 40° disk-shaped stimulus centred on this location when the eye is in its resting position. As the eyes move within their beating field during OKR, the actual, non-stationary retinal coverage extends further rostrally and caudally. For (a) red, green, and blue photoreceptors, high densities coincide with high OKR gains. (b) For ultraviolet receptors, there is no clear relationship to the OKR gain. (c) For reference, the summed total density of all receptor types combined. We did not observe a significant shift in the position-dependence of maximum OKR gain between groups of larvae at 5 dpf, 6 dpf or 7 dpf of age, consistent with the notion that retinal development is far advanced and the circuits governing OKR behaviour are stable at this developmental stage. **Figure 6–figure supplement 1.** Retinal cone densities in retinal coordinates, instead of visual field coordinates.

## Discussion

We used a spherical visual display to systematically investigate visual space anisotropies of the zebrafish optokinetic response. We show that animals react most strongly to stimuli located laterally and near the equator of their visual space. Across individuals, the OKR appears to be symmetric between both eyes, although individual animals oftentimes have a dominant eye. For small stimuli, the OKR gain depends on the size of the area in which the stimulus was presented in a logarithmic fashion. OKR to our mostly green stimuli was tuned to the higher spatial densities of red, green and blue photoreceptors in the central retina. In addition, extra-retinal processing appears to affect the preferred OKR stimulus location, as suggested by the experiments in upside-down embedded animals.

The spherical arena introduced here covers a large proportion of the surround and therefore lends itself to many other investigations of zebrafish, and of other species with limited visual acuity. In comparison to other feasible technical solutions, such as video projection setups, our spherical LED array stimulus setup provides homogeneous light and contrast across the entire stimulation area. Thereby, stimulus design is much easier because stimulus warping and conditioning becomes unnecessary. When combined with calcium imaging in a scanning microscope, the use of LED arrays provides the additional advantage that the visual stimulus can be controlled with high temporal precision, fast enough to interlace visual stimuli and line scans.

Despite the common notion that OKR is a whole-field gaze stabilisation behaviour, our results show that the OKR can be driven effectively by moving stimuli that cover only small parts of the spherical surface, e.g. a a half-maximum OKR gain is observed for a stimulus that covers 25 % of the spherical surface). Our experiment on spatial frequency dependence further demonstrates that the spatial frequency tuning of the OKR is similar across retinal locations. Since photoreceptors are not equally distributed across the retina, this result suggests that photoreceptor density is not the limiting factor for OKR performance in this frequency range.

Previous reports indicated that the zebrafish visual system is lateralised with the left eye preferentially assessing novel stimuli, while the right eye is associated with decisions to respond (45, 48). We therefore investigated whether there are consistent behavioural asymmetries for the OKR and observed almost no consistent, inter-individual asymmetries in OKR between the left and right hemispheres of the visual field, other than those induced by external conditions. Individual fish, however, show a wide and continuous range of biases towards either hemisphere.

We measured OKR gain in larvae at 5-7 days post fertilisation (dpf) of age, whereas our data on photoreceptor densities corresponds to slightly older, 7-8 dpf larvae. Owing to their rapid development, zebrafish undergo noticeable morphological changes on this timescale, but the zebrafish retina itself is known to be well developed by 5 dpf (49) and stable OKR behaviour is exhibited from then on. Crucially, we did not observe a salient age-dependent spatial shift of maximum OKR gain between our 5 dpf and 7 dpf larvae (data not shown).

The qualitative match between red cone retinal photoreceptor densities and the stimulus position driving the highest OKR gains may provide a mechanistic bottom-up explanation of the gradual differences associated with OKR. The correspondence of red photoreceptor density with the visual field map of OKR gain is consistent with the fact that our LEDs emit light at 568 nm peak power, which should have activated the red cones most. Our data is also in agreement with observations in other species, that the OKR drive is strongest when the moving stimulus covers the central visual field (10, 11, 25). In a simplistic, additive view of visual processing, increased numbers of receptors would be triggered by incident light, gradually leading to stronger activation of retinal ganglion cells and downstream circuits, eventually driving extraocular eye muscles towards higher amplitudes. Instead, or in addition, the increased resolution offered by denser distributions of photoreceptors could help reduce sensory uncertainty (and increase visual acuity). It is unclear however, how more uncertainty would lead to consistently lower OKR gains instead of a repeated switching between periods of higher and lower gains, or between OKR and other behaviours. If sensory uncertainty were indeed crucial to OKR tuning, presenting blurred or otherwise deteriorated stimuli should reduce OKR gain in disfavoured locations more strongly than those in favoured locations. It is also possible that correlations between OKR gain and photoreceptor density are entirely coincidental, as our spatial frequency tuning results for different stimulus locations had implied. Genetic zebrafish variants with altered photoreceptor distributions would thus be a valuable tool for further studies.

The pronounced increase in OKR gain for nearly lateral stimulus locations raises questions regarding the top-down behavioural significance of these directions in the natural habitat of larval zebrafish. While reduced OKR gains near the limits of the visual field might be expected, we show that gains are also reduced in the frontal binocular area, as well as in upper and lower visual field locations. Interestingly, when animals were mounted upside-down, they still prefer stimulus locations just above the equator of the environment. This result cannot be explained by shifted resting vertical eye positions in the inverted animal, which we have measured. Instead, it could potentially be explained by multimodal integration, where body orientation appears to influence the preferred OKR stimulus locations via the vestibular system (50–52). Furthermore, it seems possible that the unequal distribution of OKR gains across the visual field is related to the optic flow statistics that naturally occur in the habitats of larval zebrafish (13, 53–56). For another stabilisation behaviour of zebrafish, the optomotor response (22), we have recently shown that the underlying circuits prefer stimulus locations in the lower temporal visual field to drive forward optomotor swimming (57). Therefore, the optokinetic and the optomotor response are preferentially driven by different regions in the visual field, suggesting that they occur in response to different types of optic flow patterns in natural habitats. Both the optokinetic and the optomotor response (OKR, OMR) are thought to be mediated by the pretectum (21, 35), and we therefore hypothesize that circuits mediating OKR and OMR segregate within the pretectum and form neuronal ensembles with mostly different receptive field centre locations. Future studies on pretectal visual feature extraction in the context of naturalistic stimulus statistics are needed to establish a more complete picture of the visual pathways and computations underlying zebrafish OKR, OMR and other visually mediated behaviours.

## Methods

### Animal experiments

Animal experiments were performed in accordance with licenses granted by local government authorities (Regierungspräsidium Tübingen) in accordance with German federal law and Baden-Württemberg state law. Approval of this license followed consultation of both in-house animal welfare officers and an external ethics board appointed by the local government. We used *mitfa*−/− animals (5-7 dpf) for the experiments, because this strain lacks skin pigmentation that could interfere with eye tracking.

### Coordinate systems and conventions

To remain consistent with the conventions adopted to describe stimuli and eye positions in previous publications, we adopted an East-North-Up, or ENU, geographic coordinate system. In this system, all positions are relative to the fish itself, and expressed as azimuth (horizontal angle, with positive values to the right of the fish), elevation (vertical angle, with positive values above the fish), and radius (or distance to the fish). The point directly in front of the fish (at the rostrum) is located at [0°, 0°] azimuth and elevation. Azimuth angles cover the range [−180°, 180°] and elevation angles [−90°, 90°]. Azimuth sign is opposite to the conventional mathematical notation of angles when looking top-down onto the fish. Supplementary materials provide a detailed description of the coordinate systems used, including transformations between Cartesian and geographic coordinate systems **(Supplementary File 2)**.

### Design of the spherical arena

#### Geometric design of the arena

The overall layout of the spherical arena was optimised to contain the near maximum number of LED tiles that can be driven by our hardware controllers (232 out of a possible 240), and arrange them with minimal gaps in between. Also, care was taken to leave sufficient gaps near the top and bottom poles to insert the optical pathway used to illuminate and record fish behaviour. A further 8 LED tiles could be included as optional covers for the top and bottom poles, bringing the total number to 240 out of 240 possible. A detailed walkthrough of the mathematical planning is found in the supplementary material **(Supplementary File 2)**.

#### Arena elements

The arena consists of a 3D-printed structural scaffold; green light emitting LED tiles (Kingbright TA08-81CGKWA, 20×20 mm each, peak power at 568 nm) hot-glued to the scaffold and connected by cable to a set of circuit boards with hardware controllers **(Figure 1–figure supplement 1d)**; 8×8 individual LEDs contained in each tile **(Figure 1f)**; a nearly spherical glass bulb filled with water, into which the immobilised larvae are inserted (**Figure 1–figure supplement 1c**, middle); a metal rotation mount attached to the scaffold “keel” of the arena (**Figure 1–figure supplement 1c**, right), holding the glass bulb in place and allowing corrections of pitch and roll angles; the optical pathway with an infrared light source to illuminate the fish from below **(Figure 1–figure supplement 1b)**, and an infrared-sensitive USB camera for video recording of the transmission image **(Figure 1–figure supplement 1d)**.

#### Electronics and circuit design

To provide hardware control to the LEDs, we used circuit boards designs and C controller code provided by Alexander Borst (MPI of Neurobiology, Martinsried), Väinö Haikala and Dierk Reiff (University of Freiburg) (58). The electronic and software architecture of stimulus control has originally been designed by Reiser et al. (1) (also see the available documentation for a different version of the CAD and code at https://bitbucket.org/mreiser/panels/wiki/Home). Any custom circuit board design and code could be substituted for these, and alternative solutions exist, e.g., in Drosophila vision research (59). At the front end, these electronics control the 8×8 LED matrices, which are multiplexed in time to allow control of individual LEDs with just 8 input and 8 output pins.

#### Optical pathway, illumination and video recording

A high-power infrared LED was placed outside the stimulus arena and its light diffused by a sheet of milk glass and then guided towards the fish through the top hole of the arena **(Figure 1–figure supplement 1b, Figure 1–figure supplement 1d)**. Non-absorbed IR light exits through the bottom hole, where it is focused onto an IR-sensitive camera. Between the arena and the proximal lens, a neutral density filter (NE13B, Thorlabs, ND 1.3) was inserted half-way (off-axis) into the optic pathway using an optical filter slider (CFH2/M, Thorlabs, positioned in about 5 cm distance of the camera CCD chip) to improve image contrast (oblique detection). We used the 840nm, 125 degree IR emitter Roschwege Star-IR840-01-00-00 (procured via Conrad Electronic GmbH as item 491118-62) in custom casing, lenses LB1309 and LB1374, mirror PF20-03-P01 (ThorLabs GmbH), and IR-sensitive camera DMK23U618 (TheImagingSource GmbH). Approximate distances between elements are 14.5cm (IR source to first lens), 12cm (first lens to centre of glass bulb), 22cm (bulb centre to mirror centre), 8.5cm (mirror centre to second lens), 28.5 cm (second lens to camera objective).

#### Fish mounting device

Larvae were mounted inside a custom-built glass bulb **(Figure 1– figure supplement 1c, middle)**. Its nearly spherical shape minimises reflection and refraction at the glass surface. It was filled with E3 solution, so there was no liquid-to-air boundary distorting visual stimuli. Through an opening on one side, we inserted a glass rod, on the tip of which we immobilise the larva in agarose gel (see description of the embedding procedure below). The fish was mounted in such a way that the head protruded the tip of the narrow triangular glass stage, which ensured that visual stimuli are virtually unobstructed by the glass triangle on their way to the eyes **(Figure 1–figure supplement 1c, left)**. The entire glass structure was held at the centre of the spherical arena by metal parts attached to the arena scaffold itself **(Figure 1–figure supplement 1c, right)**. Care was taken to remove air bubbles and completely fill the glass bulb with E3 medium.

#### Computer-assisted design and 3D printing

To arrange the square LED tiles across a nearly spherical surface, we 3D-printed a structural scaffold or “skeleton”, consisting of a reinforced prime meridian major circle (“keel”) and several lighter minor circles of latitude **(Figure 1g)**. Available hardware controllers allow for up to 240 LED matrices in parallel, so we chose the exact size of the scaffold (106.5 mm in diameter) to hold as many of these as possible while minimising gaps in between. As individual LEDs are arranged in a rectangular pattern on each of the flat LED tiles, and stimuli defined by true meridians (arcs from pole to pole, or straight vertical lines in Mercator projection), pixelation of the stimulus is inevitable, and stimulus edges become increasing stair-shaped near the poles. Because of the poor visual acuity of zebrafish larvae (see **Supplementary File 2**), this should not affect OKR behaviour. Our design further includes two holes necessary for behavioural recordings and two-photon imaging, located at the North and South poles of the sphere. We placed the largest elements of the structural scaffold behind the zebrafish **(Figure 1–figure supplement 1d)**. Given the ~160° azimuth coverage per eye in combination with a slight eye convergence at rest, this minimises the loss of useful stimulation area.

We printed all structures out of polylactide (PLA) filament using an Ultimaker 2 printer (Ultimaker B.V.). Parts were assembled using a hot glue gun.

### Visual field coverage

We can estimate the fraction of the visual field effectively covered by LEDs based on a projection of LED tiles onto a unit sphere. The area *A* of a surface segment delimited by the projection of the edges of a single tile onto the sphere centre is given by

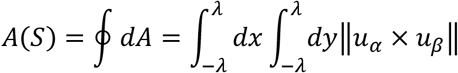

where *u*_*α*_ and *u*_*β*_ are the Cartesian unit vectors spanning the tile itself and (±λ, ±λ) is the Cartesian position of the four edges of another rectangle. This smaller rectangle is the straight projection of the sphere segment onto the tile,

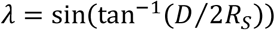

where *R*_*S*_ = 106.5 *mm* is the sphere radius and *D* = 21 *mm* is the length of the edges of the tile. Summing over the number of tiles included in the arena, the equations above can be used to estimate the total coverage of the sphere by its square LED tiles to around 66.5% of the surface area. Using this strict estimate, the small gaps in between LED arrays are counted as not covered, even though we successfully demonstrated that they are small enough not to affect OKR performance, likely due to the low visual acuity of zebrafish larvae. A more meaningful estimate of coverage must take these results into account **(Supplementary File 1)**, and in fact reveals that stimuli presented with our LEDs effectively cover 85.6% of all possible directions. In core parts of the visual field, coverage exceeds 90%.

### Stimulus design

We designed visual stimuli, transformed them to geographical coordinates, and mapped them onto the physical positions of each individual LED with custom MATLAB software. We have made this code available for free under a Creative Commons NC-BY-SA 4.0 license **(Supplementary Code 1)**. The mapped stimulus was then uploaded to the hardware controllers using custom-built C code originally developed by Väinö Haikala.

To investigate OKR gain dependence on stimulus location, we chose to present stimuli centred on 36 different locations distributed nearly equidistantly across the spherical arena, as well as symmetrically distributed between the left and right, upper and lower, front and rear hemispheres **(Figure 3a)**. These positions were determined numerically: First, we populated one eighth of the sphere surface by placing one stimulus centre at a fixed location at the intersection of the equator and the most lateral meridian (90 degrees azimuth, 0 degrees elevation), constraining two more stimulus centres to move along this lateral meridian (90 degrees azimuth, initially random positive elevation), constraining yet another stimulus centre to move along the equator (initially random positive azimuth, 0 degrees elevation), and allowing three more stimulus centre to move freely across the surface of this eighth of the sphere (initially random positive azimuth and elevation), for a total of 7 positions. Second, we placed additional stimulus centres onto all 29 positions that were mirror-symmetric to the initial 7, with mirror planes placed between the six hemispheres listed above. We then simulated interactions between all 38 stimulus centres akin to electromagnetic repulsion, until a stable pattern emerged. Resulting coordinate values were rounded for convenience **(Supplementary Code 2, Video 1)**.

### Embedding procedure

To immobilise fish on the glass tip inside the sphere, we developed a novel embedding method. A cast of the glass triangle (and of the glass rod on which it is mounted) was made by placing it inside a Petri dish, which was then filled with a heated 2% agarose solution. After agarose cooled down and polymerised, agarose within a few millimetres of the tip of the glass triangle was manually removed, before removing the triangle itself. The resulting cast was stored in a refrigerator and then used to hold the glass triangle during all subsequent embedding procedures, limiting the freedom of movement of the larva to be embedded. The triangle was stored separately at room temperature. Before each embedding, we coated the glass triangle with polylysine and dried it overnight in an incubator at 29 degrees Celsius to increase the subsequent adhesion of agarose. We then returned the glass triangle into its cast, and constructed a tight, 2 mm high circular barrier around its tip using pieces of congealed agarose. A larva was picked up with as little water as possible using a glass pipette and very briefly placed inside 1 ml of 1.6% low-melting agarose solution at 37 degrees Celsius. Using the same pipette, the larvae was then transferred onto the glass triangle along with the entire agarose. After the larva had been placed a few millimetres away from the tip of the glass triangle, the orientation of the animal could be manipulated with custom-made platinum wire tools without touching its body, as previously described (60). Before the agarose congeals, swimming motions of the animal were exploited to guide it towards the tip and ensure an upright posture. The final position of the fish was chosen as such that its eyes are aligned with the axis of the glass rod, its body is upright without any rotation, and its head protrudes forward from the tip of the glass triangle, maximising the fraction of its field of view unobstructed by glass elements. The agarose was left to congeal, and the Petri dish was filled with in E3 solution. The freshly congealed agarose surrounding the glass triangle was then removed using additional, flattened platinum wire tools, once again separating the glass triangle from the cast. Using the same tools, we finally cut triangular holes into the remaining agarose to completely free both eyes. To ensure free movement of both eyes, we confirmed the presence of large and even optokinetic eye movements using a striped paper drum before the experiment.

We then pick up the glass triangle by the glass rod attached to it, cut off any remaining agarose detritus, and place it inside the E3-filled glass bulb. No air remained in the bulb, and no pieces of detritus were introduced into the bulb, as these would accumulate near the top and bottom of the bulb, respectively, interfering with the optical pathway and thus reduce image quality.

### Data analysis

Video images of behaving zebrafish larvae were processed in real time using a precursor of the *ZebEyeTrack* software (40), available from www.zebeyetrack.com. The resulting traces of angular eye position were combined with analogue output signals from the hardware controllers of the spherical arena to match eye movement to the various stimulus phases. This was achieved using custom-built MATLAB software, which is freely available under a Creative Commons NC-BY-SA 4.0 license **(Supplementary Code 3)**.

Data was then analysed further by detecting and removing saccades and fitting a piece-wise sinusoidal function to the eye position traces. The parameters of the fit were then compared to the parameters of the equally sinusoidally changing angular positions of the stimulus. For each fish, eye, and stimulus phase, the ratio between the amplitude of the fit to eye position and the amplitude of stimulus position represents one value of the gain of the optokinetic response.

For each interval between two subsequent saccades, or inter-saccade-interval (ISI), the fit function to the eye position data is defined by

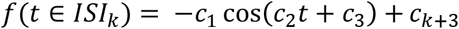

Here, *t* are the time stamps of data points falling within the *k*-th ISI, *c*_1_, *c*_2_ and *c*_3_ are the amplitude, frequency and phase shift of oscillation across all ISIs, and *c*_*k*+3_ is a different constant offset within each ISI, which corrects for the eye position offsets brought about by each saccade. The best fit value *c*_1_ was taken as an approximation of the amplitude *a*_*E*_ of eye movement, *a*_*E*_ ≈ *c*_1_. The process of cropping saccades from the raw data and fitting a sinusoid to the remaining raw data is demonstrated in **Figure 2–Figure supplement 1**.

The OKR gain *g* is a common measure of visuomotor function. It is defined as the ratio between the amplitude *a*_*E*_ of eye movement and the amplitude *a*_*S*_ of the visual stimulus evoking eye movement,

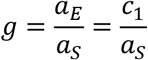

In other words, OKR gain indicates the degree to which zebrafish larvae track a given visual stimulus. For each eye, a single gain value per stimulus phase is computed. While a value of 1 would indicate a “perfect” match between eye movement and stimulus motion, zebrafish larvae at 5 dpf often exhibit much lower OKR gains (33). While highest gains are obtained for very slowly moving stimuli, in our experiments, we chose higher stimulus velocities. Although these velocities are only tracked with small gains, the absolute velocities of the eyes are high, which allowed us to collect data with high signal-to-noise levels and reduce the needed recording time.

To rule out asymmetries induced by the arena itself or by its surroundings, we recorded two sets of stimulus-position-dependence data, one with the arena in its original configuration, and another with the arena rotated by 180 degrees **(Figure 4a-b)**. Each set contained data from multiple larvae, and with at least 2 separate presentations of each stimulus position. For each stimulus position, and separately for both sets of data, we computed the median OKR gain across fish and stimulus repetitions. We then averaged between the two datasets, yielding a single OKR gain value per stimulus position. As asymmetries are less crucial when studying stimulus frequency and size **(Figure 5)**, we did not repeat those with a rotated arena, and could thus omit the final step of the analysis.

### Von Mises-Fisher fits to data

Based on the assumption that OKR position tuning could be normally distributed with respect to each angle, OKR gain would be approximated by a two-dimensional, circular von Mises-Fisher function centred on the preferred stimulus location. Because the eyes are yoked, the OKR gain of one eye will be high around its own preferred position, as well as around the preferred position of the contralateral eye. To account for this, we fit the sum of two independent von Mises-Fisher functions to our OKR gain data:

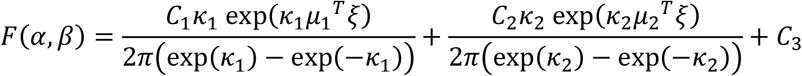

Here, *ξ* is the Cartesian coordinate vector of a point on the sphere surface, and corresponds to the geographic coordinates azimuth *α* and elevation *β*. *μ*_1_ and *μ*_2_ are Cartesian coordinate vectors pointing to the centre of the two distributions, and *κ*_1_ and *κ*_2_ express their respective concentrations, or narrowness. The parameters *μ*_j_, *κ*_j_, the amplitudes *C*_1_, *C*_2_ and the offset *C*_3_ are fit numerically.

**Figure 3** and **Figure 4e-h** show the best von-Mises-Fisher fits to data as coloured sphere surfaces, whereas the individual data points are shown as small coloured circles in **Figure 3**. All results shown in **Figure 3** and **Figure 4e-h** pertain to a single, constant size of cropping disks, and the colour of any point across the sphere surface represents the expected value of OKR gain if another stimulus were shown with a cropping disk centered on this location.

### Yoking index, asymmetry and mathematical modelling

To quantify asymmetries in the gain between left and right, stimulated and unstimulated eyes, we introduce the yoking index

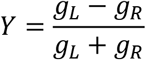

Here, *g*_*L*_ and *g*_*R*_ are the OKR gains of the left eye and right eye, measured during the same stimulus phase. Depending on stimulus phase, only the left eye, only the right eye or both eyes may have been stimulated. If the yoking index is positive, the left eye responded more strongly than the right eye; if it is negative, the amplitude of right eye movement was larger. An index of zero indicates “perfect yoking”, i.e. identical amplitudes for both eyes.

In addition, we define a “bias” index to capture innate or induced asymmetries between responses to stimuli presented within the left or right hemisphere of the visual field,

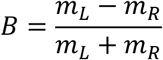

Here, *m*_*L*_ and *m*_*R*_ are the medians of OKR gains after pooling across either all left-side or all right-side stimulus types (D1-D19 and D20-D38, respectively). Several sources of asymmetry contribute to *B*: (1) arena- or environment-related differences in stimulus perception, constant across individuals; (2) a biologically encoded preference for one of the two eyes, constant across individuals; (3) inter-individual differences between the eyes, constant across stimulus phases for each individual; (4) other sources of variability unaccounted for, and approximated as a noise term *η*. We hypothesise that the overall asymmetry observed for each larva *k* is given by a simple linear combination of these contributions,

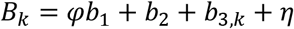

The parameter *φ* is 1 for the default arena setup, and −1 during control experiments with a horizontally flipped arena setup. To determine *b*_1_, *b*_2_ and *b*_3_, we fit this system of equations by multivariate linear regression to experimentally observed bias indices. The system is initially underdetermined, as it contains *n* + 2 coefficients for every *n* fish observed. However, if we assume that individual biases average out across the population, we can determine the population-wide coefficients *b*_1_ and *b*_2_ by setting aside the individual *b*_3,*k*_ for a first regression. To determine how far each individual deviates from the rest of the population, we then substitute their best regression values of *b*_1_ and *b*_2_ into the full equation, and perform a second regression for the remaining *b*_3,*k*_.

## Acknowledgements

We thank Väinö Haikala and Dierk F. Reiff (University of Freiburg) for sharing their code and design for hardware controllers, Alexander Borst (MPI Neurobiology, Martinsried) for providing the LED panel board design (these designs are based on original precursory versions designed by Reiser et al. (1), Thomas Nieß (glassblowing workshop, University of Tübingen) and Klaus Vollmer (precision mechanics workshop, University of Tübingen) for technical support, Prudenter-Agas (Hamburg, Germany) for generating glass bulb illustrations, Andre Maia Chagas for help with 3D printing procedures, and Jan Benda (University of Tübingen) for discussions on visual acuity.

## Supplementary information

### Supplementary Code

**Supplementary Code 1.** (S1Code.zip) MATLAB code to design visual stimuli, convert them to geographic coordinates, and map them onto the actual position of individual LEDs.

**Supplementary Code 2.** (S2Code.zip) MATLAB code to numerically identify a distribution of nearly equidistant stimulus centres that is symmetric between the left and right, upper and lower, as well as front and rear hemispheres.

**Supplementary Code 3.** (S3Code.zip) MATLAB code to read raw eye traces, identify individual stimulus phases, detect and remove saccades, compute piece-wise fits to cropped and pre-processed raw data, and return OKR gains.

**Supplementary Code 4.** (S4Code.zip) MATLAB code to recreate the results figures from data, especially **Figure 3, Figure 5, Figure 6**.

**Supplementary Code 5.** (S5Code.zip) MATLAB code to assess the significance of differences between the best fits to data for fish embedded upright or upside-down. This code requires access to the raw data repository.

### Supplementary Manuals

**Supplementary File 1.** (S1Text.pdf) Coverage of visual field by stimulus arena

**Supplementary File 2.** (S2Text.pdf) Spatial resolution and visual acuity

**Supplementary File 3.** (S3Text.pdf) Mathematical appendix on the design of and stimulus mapping onto the spherical arena, written as a manual.

### Supplementary Figures

**Figure 1–figure supplement 1.**
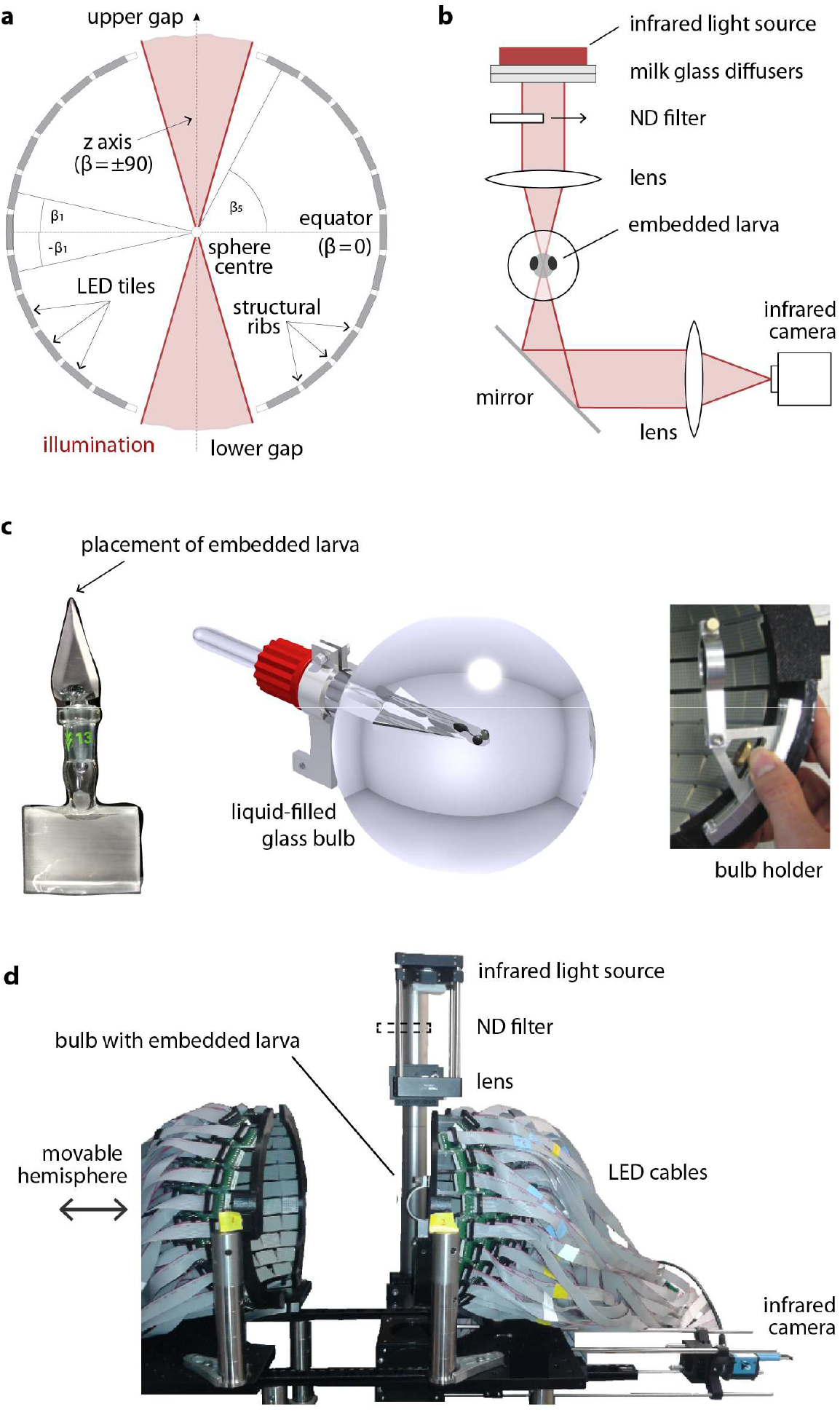
A spherical LED arena to present visual stimuli across the visual field. (a) LED tiles are arranged in ribbons parallel to the equator, and glued in between structural ribs. Gaps at the top and bottom pole of the sphere allow coupling in an optical pathway for infrared illumination and subsequent video recording of eye movement. The fish is placed at the centre of the sphere, facing the observer. See **Supplementary File 3** and **Supplementary File 4** for more on the angles β_k_. (b) Optical pathway for eye movement tracking. (c) To minimise obstruction and refraction, the zebrafish larva is immobilised on the tip of a glass triangle (left) using agarose, which is then inserted into the centre of a spherical glass bulb (middle). This bulb is then mounted into a metal holder (right) and thus placed at the centre of the sphere. (d) Image of the two hemispheres and the camera setup. One hemisphere is mounted on a rail to allow opening and closing the arena.

**Figure 2–figure supplement 1.**
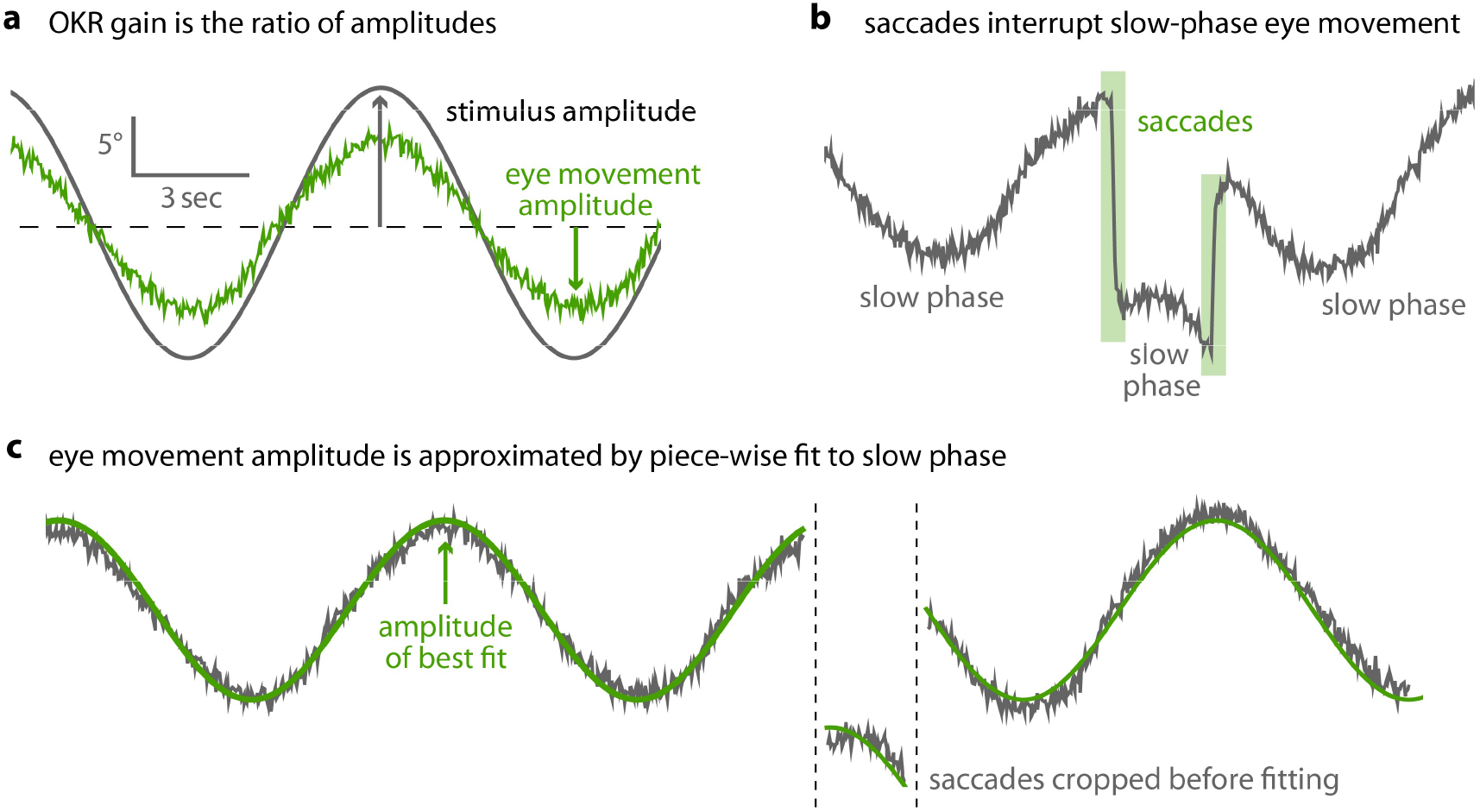
Saccades are cropped before OKR gain is measured. (a) As shown in **Figure 2b**, optokinetic gain is the ratio between the amplitudes of eye movement and stimulus movement. To extract these, raw eye traces must be processed. (b) OKR eye movements consist of a slow phase, gradually tracking stimulus motion, and intermittent saccades. (c) After pre-processing data to detect and remove saccades, we fit a piece-wise sinusoidal function with a single amplitude to the remaining slow-phase eye traces. The amplitude of the best fit determines OKR gain.

**Figure 2–figure supplement 2.**
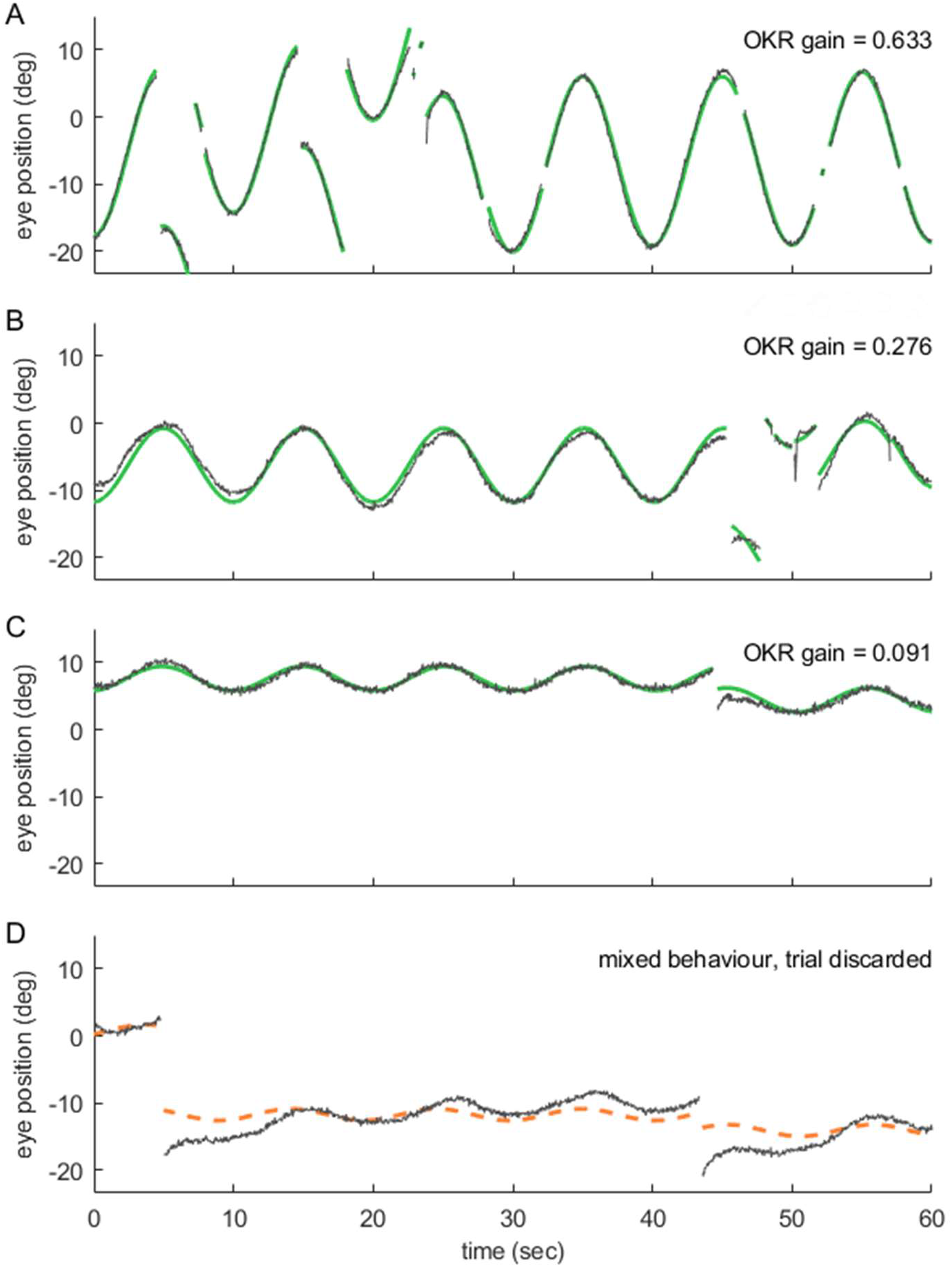
Even at remote stimulus locations, fish exhibit reliable OKR behaviour; trials with mixed behaviour are excluded. (a-d) show 60-second samples of eye position traces obtained during visual stimulation (dark gray traces). Saccades were cropped, as demonstrated in **Figure 2-figure supplement 1**. Positive angles correspond to more rightward eye positions. Coloured traces show best fit to data (see **Methods**), along with the resulting OKR gain. Fish were presented with either (a) whole field visual stimuli, or (b-d) smaller disk-shaped stimuli, as in **Figure 3**. Whole-field stimulation elicits reliable OKR, even though high stimulus frequencies keep its gain below 1. (b) The same is true for disk stimulus in a preferred region of the visual space (stimulus D7, azimuth −90°, elevation +39°, i.e. lateral and dorsal to the fish), although the smaller stimulus evokes a smaller OKR gain. (c) Disk stimuli presented in a disfavoured part of the visual field (stimulus D3, azimuth −90°, elevation +74°, almost dorsal to the fish) evoke significantly lower gains, but still reliable OKR behaviour. Data in (a,c) correspond to trials shown in **Figure 2c**. (d) As it is possible to encounter behaviours other than OKR, or in addition to OKR, we inspected all trials before data analysis. Trials in which no pure OKR was present, such as the one shown here, were excluded (stimulus D3, same as in (c)).

**Figure 3–figure supplement 1.**
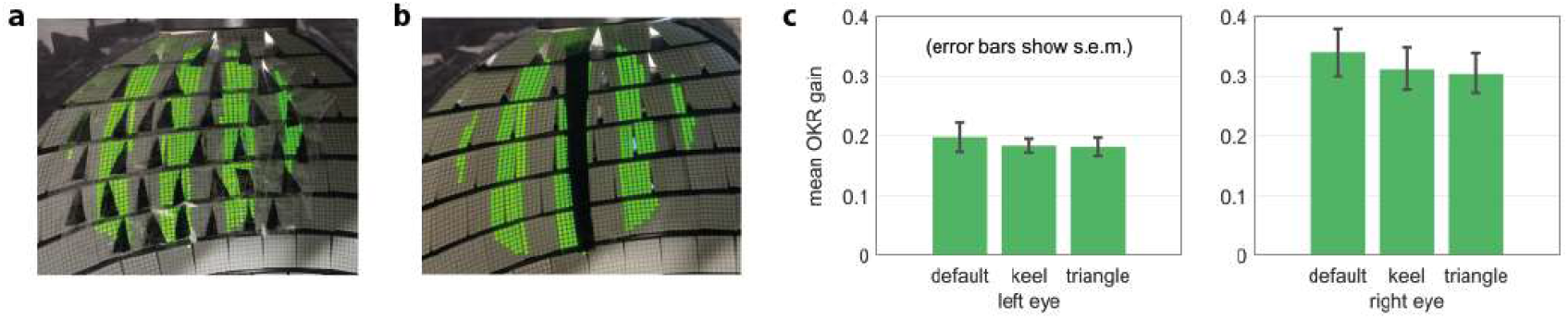
Gaps in the arena do not bias OKR behaviour. (a) Artificial triangular holes setup. (b) Artificial keel setup. (c) Neither triangular nor elongated gaps result in significantly different OKR gains.

**Figure 3–figure supplement 2.**
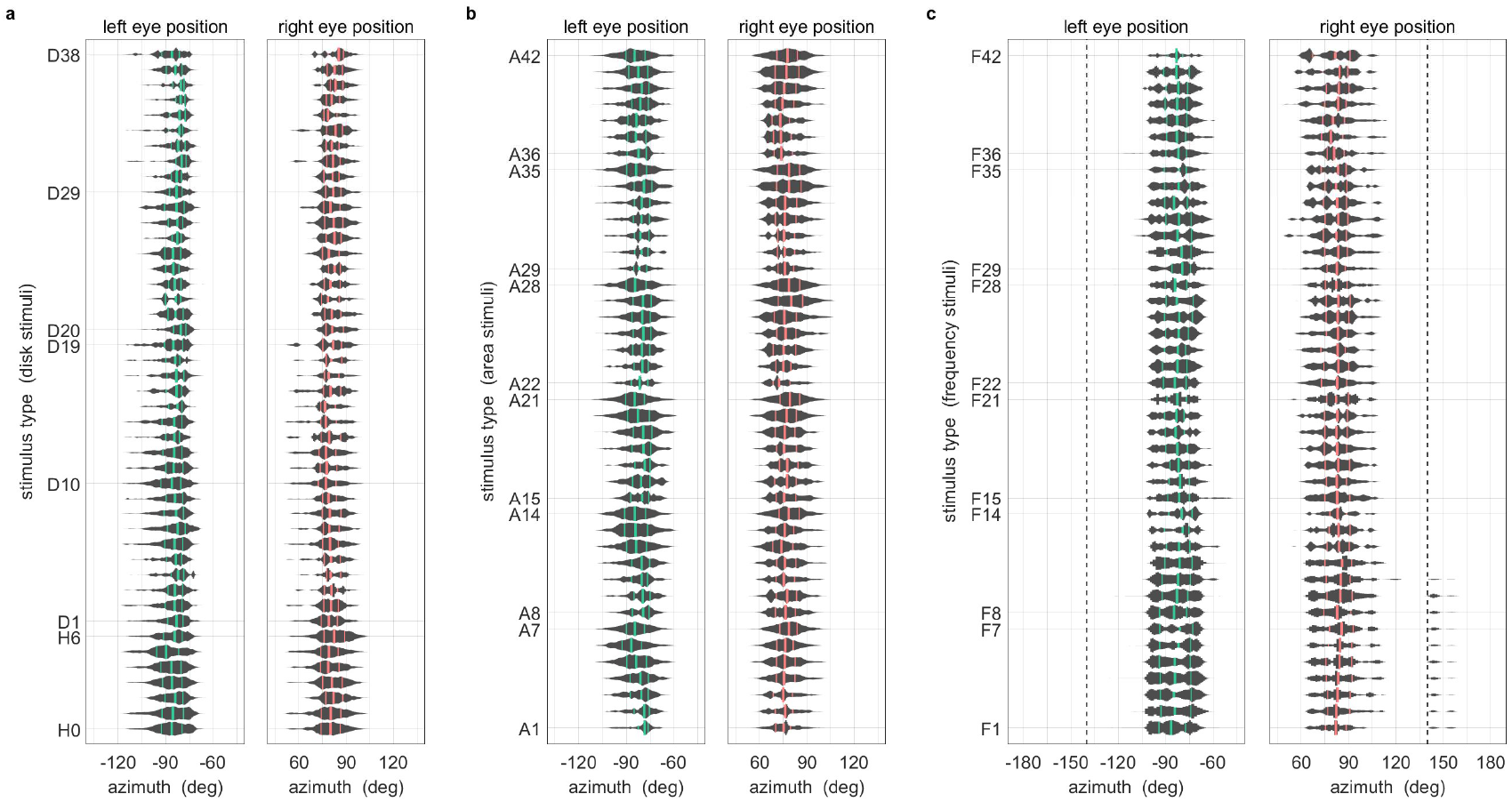
During OKR, the beating field and average eye position are independent of stimulus location. All data were pooled across fish and trials. One gain value was computed per stimulus presentation. Violin plots show distribution of mean horizontal eye positions across the pooled data; vertical lines indicate 25th percentile, median and 75th percentile of the distribution. Positions are those during presentation of stimulus types (a) D1 to D38 shown in **Figure 3a**, (b) A1 to A42 shown in **Figure 5b**, (c) F1 to F42 shown in **Figure 5a**. Dashed lines in (c) represent axis limits of (a,b).

**Figure 3–figure supplement 3.**
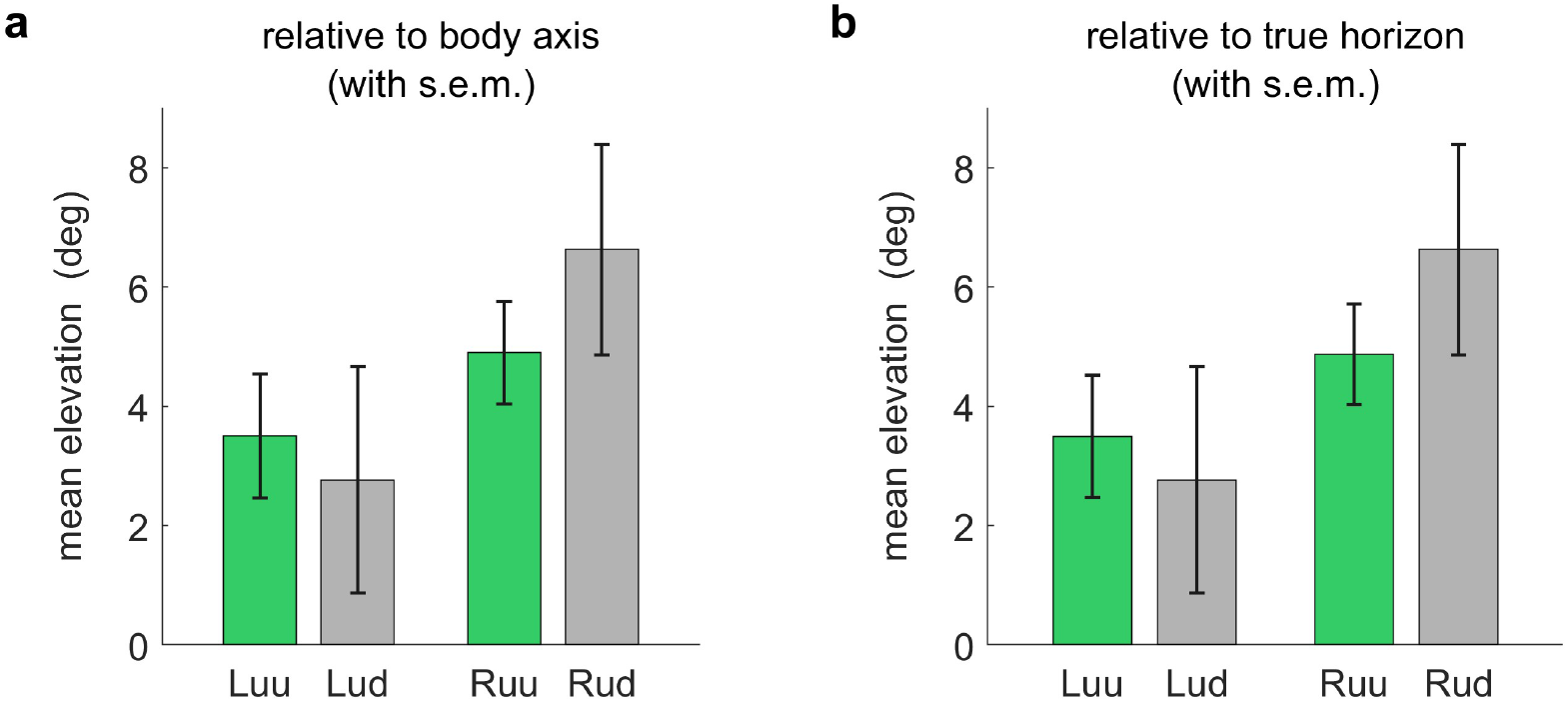
Vertical eye position under upright and upside-down embedding. Larvae were embedded in agarose with their eyes free, and placed under a microscope. Using an additional mirror, we recorded image time series from the front, i.e., looking along the anterior-posterior axis. Vertical eye position was determined geometrically for each individual frame. Because some larvae were embedded in such a way that mediolateral axis was not entirely aligned with the true horizon of the environment, we measured vertical eye position (a) relative to both the mediolateral body axis, and (b) the mediolateral body axis, but corrected by its offset from the true horizon. Required corrections were minimal. More importantly, there were no significant differences between the vertical eye positions of larvae embedded upside-up (uu) or upside-down (ud), neither for their left (L) or right eyes (R). Bars indicate mean after pooling across all frames and individuals, error bars show corresponding standard error of the mean (s.e.m.). On all panels, positive angles indicate dorsalward eye positions. In summary, larval eyes were almost always inclined towards the dorsum, irrespective of the direction of embedding. The fish do not appear to compensate for their orientation with respect to the gravitational axis.

**Figure 3–figure supplement 4.**
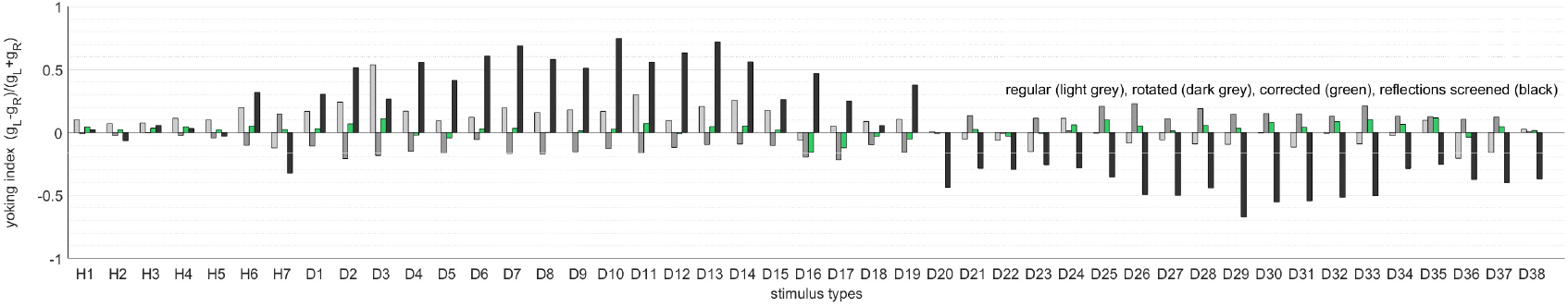
Yoking indices are biased by reflections within the arena. Yoking indices were computed for experiments using the regular setup as in **Figure 4a** (light grey), with a rotated arena as in **Figure 4b** (dark grey), corrected for experimental asymmetries as in **Figure 3** (green), and with one side of the glass bulb, contralateral to the stimulus centre, painted black (black). Yoking indices from most experiments are close to zero, indicating similar OKR gains for both eyes regardless of stimulus location. In contrast, yoking indices from the latter control experiment differ markedly from zero, indicating significantly weaker responses by the respectively unstimulated eye. This finding points to reflections at the air-glass interface being visible to the purportedly “unstimulated” eye.

**Figure 3–figure supplement 5.**
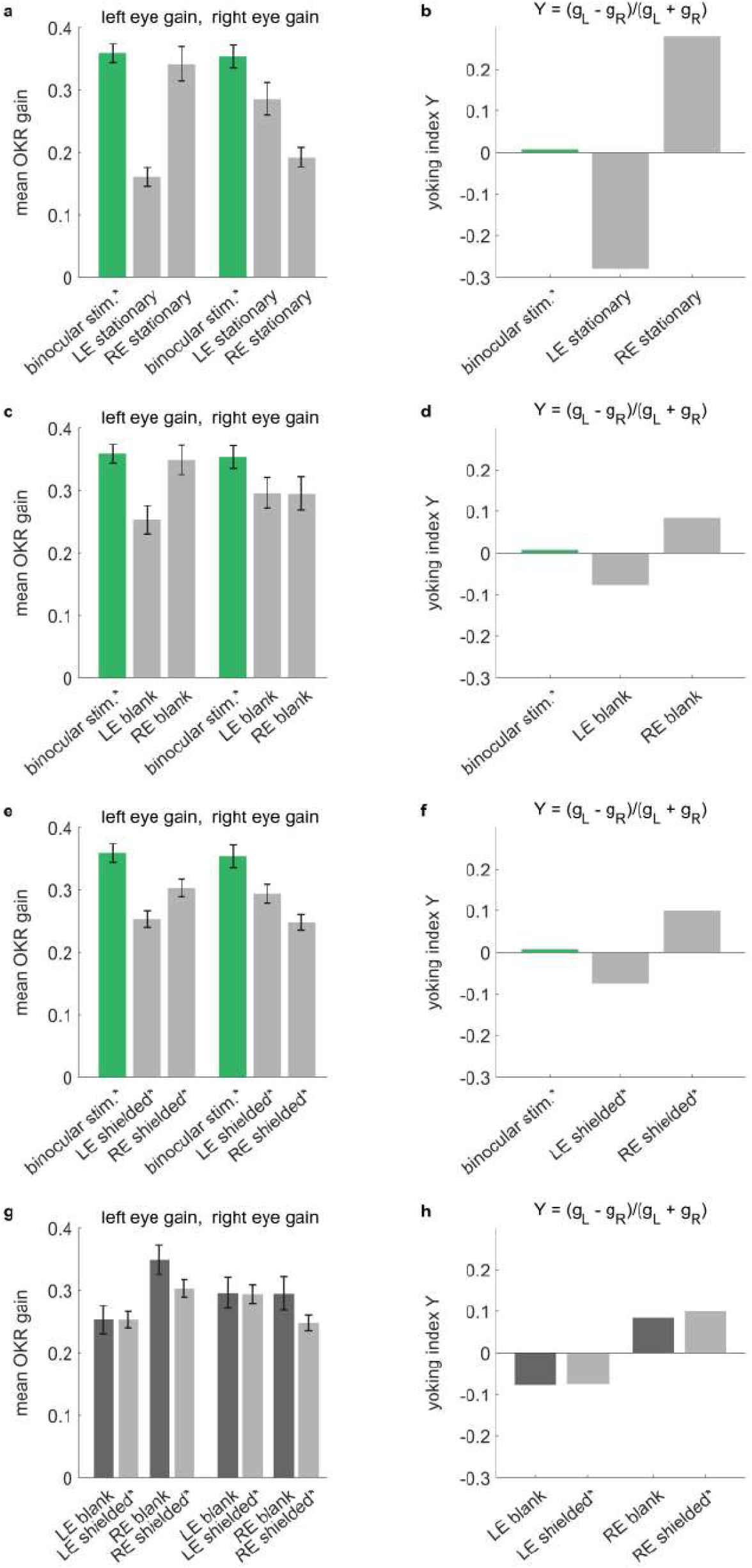
Across different types of arenas, stimulus reflections affect perceived yoking. Control experiments conducted in a rectangular stimulus arena. OKR-inducing gratings were shown on all four stimulus screens surrounding the larvae, while additional elements were introduced around either the left eye (LE), the right eye (RE) or neither. Specifically, selected eyes were (a,b) shown stationary stimuli of the same frequency and contrast as the moving stimuli, (c,d) shown a blank white surface, or (e,f) shielded with a fully opaque cover. (a,c,e,g) Bars indicate mean OKR gains, and error bars show standard error of the mean. (b,d,f,h) Yoking indices are near zero when both eyes move with identical amplitudes and positive when left eye amplitude exceeds that of the right eye (see **Methods**). (a,b) In the presence of two conflicting stimuli (moving vs. stationary), yoking between the eyes is reduced by almost half, confirming that the unstimulated eye is yoked to the stimulated eye, albeit with a lower OKR gain. (c,d) When there is no conflicting stimulus, yoking drives OKR of the contralateral eyes, albeit with a lower amplitude as if both eyes were stimulated directly with identical stimulus, (e,f) which is equally true in the presence of shielding. (g,h) To assess the effect of reflections on the difference between directly stimulated and purportedly stimulated eyes, we compare blank stimuli (as in c-d, which could diffusely reflect light) to fully shielded eyes (as in e-f, where no reflections should occur). There are no significant differences, indicating that the larger effect of reflections observed in our spherical arena **(Figure 3–figure supplement 4)** may be caused specifically by sharp reflections of the stimulus patterns at the air-water interface, instead of more diffuse reflections of light across the background. *Two rectangular arena setups were used for the control experiments. Asterisks indicate data obtained from the second setup, for which balanced illumination was explicitly confirmed via diode photodetector. Data from n=22 fish for initial setup and n=10 fish for second setup.

**Figure 4–figure supplement 1.**
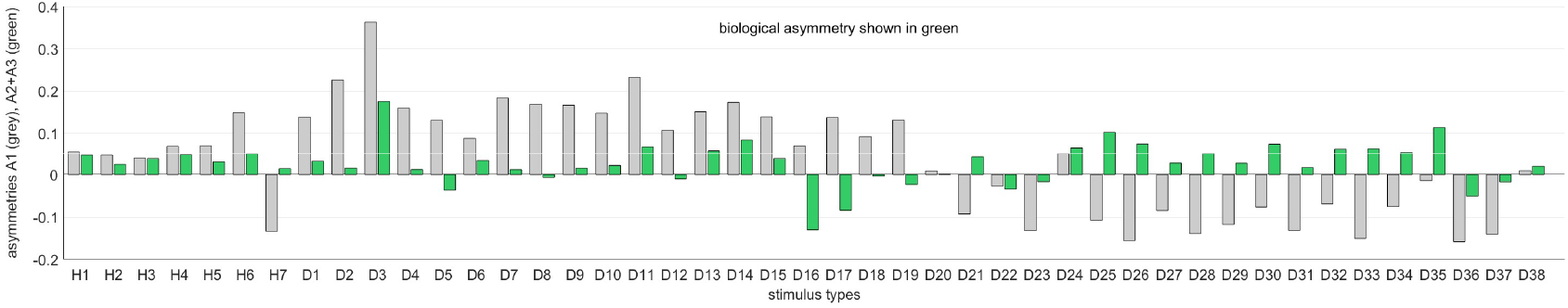
Asymmetries between left and right eye are strongly affected by the environment. Differences between the OKR gains of directly stimulated left eyes and directly stimulated right eyes can be explained by a linear combination of biological and environmental factors *b*_*k*_, e.g., biases of individual animals or asymmetries of the stimulus arena (see **Methods**). Comparing data from the regular and rotated setups **(Figure 4e-f)**, as well as data from fish immobilised upside-down **(Figure 4g)**, including additional stimuli covering entire hemispheres **(Supplementary File 7)**, we can infer the underlying *b*_*k*_ via multivariate regression of our linear model. We find that individual biases (grey) vary strongly from fish to fish, and are broadly distributed from left to right. There are some constant biases across fish (green), both towards the left side of their visual field (*b*_2_) and towards one of the two LED hemispheres (*b*_1_); however, these biases are small and, given the large variability of individual biases, might be a result of the limited number of fish studied.

**Figure 5–figure supplement 1.**
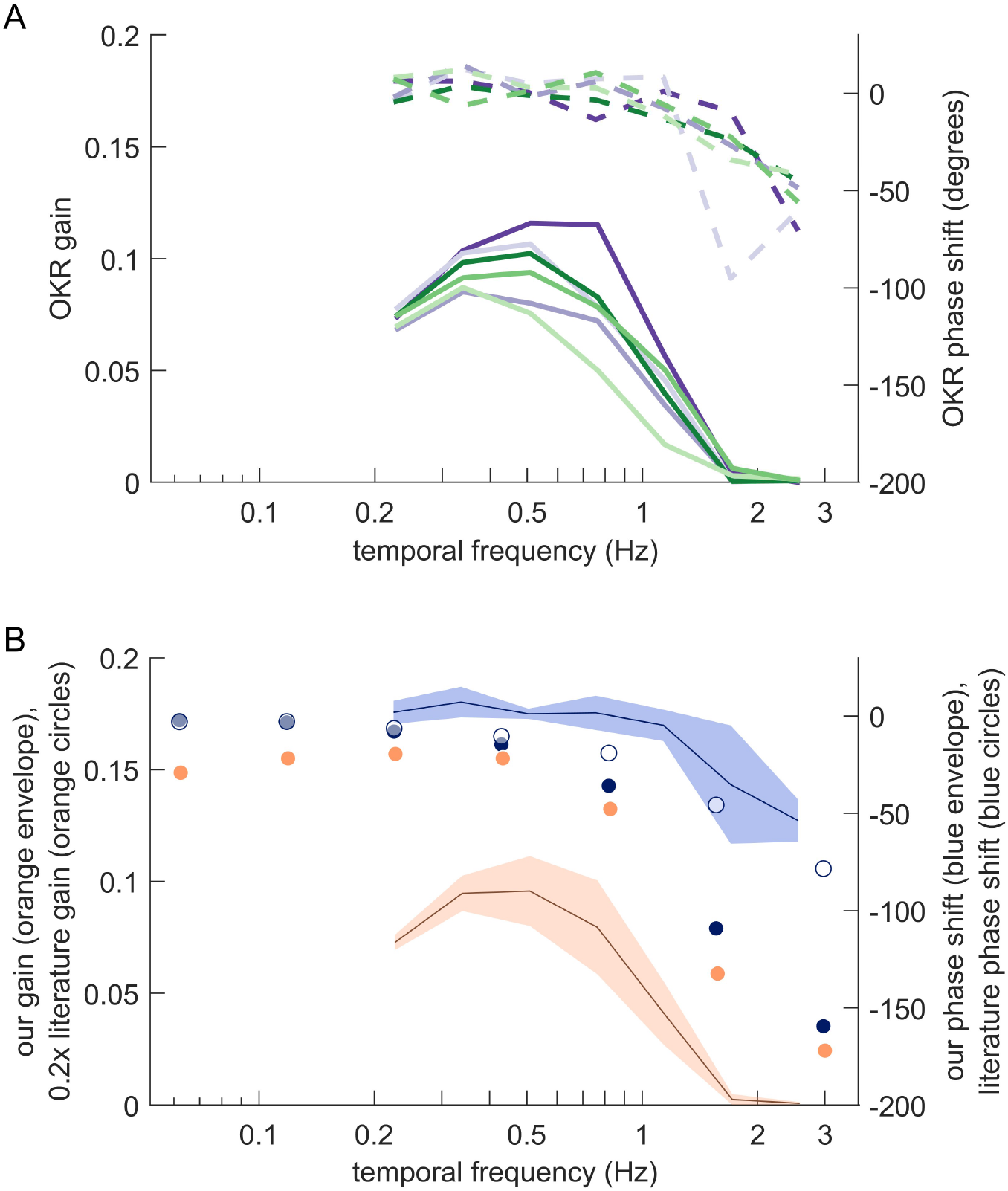
Magnitude and phase shift of eye movements at different frequencies resemble those previously observed for zebrafish OKR. (A) Bode plots showing OKR gain (solid lines) and phase shift relative to the stimulus (dashed lines). Gains and phase shifts decrease as temporal frequency increases. Each line represents the mean across trials for a specific stimulus location, pooled across all fish, all trials and both eyes. Colours match the stimulus locations identified in **Figure 5c**. (B) Our observations qualitatively, but not quantitatively, match those reported for traditional full-field stimulus paradigms. Solid blue circles represent OKR phase shifts for approximately 6.4 dpf zebrafish larvae reported by (42). The blue line represents our own mean phase shift across all stimulus locations shown in (A), and the light blue envelope shows the standard deviation across locations. Interestingly, our phase shifts are more in line with those reported for older fish (approximately 33.8dpf larvae, open blue circles) in (42). Solid orange circles show scaled OKR gain for approximately 6.4dpf zebrafish larvae, as reported by (42). Direct comparison to our gain data could be misleading because of the smaller size and higher velocity of our stimuli. Based on our results reported in **Figure 5e**, the size difference alone should account for a factor of about 5. Orange circles thus show the literature gain data scaled 0.2x. The black line and orange envelope represent our mean OKR gain and standard deviation across all stimulus locations shown in (A). Quantitative changes are consistent with the notion that smaller and faster stimuli evoke equally reliable, but lower-amplitude OKR behaviour (cf. **Figure 2c**).

**Figure 6–figure supplement 1.**
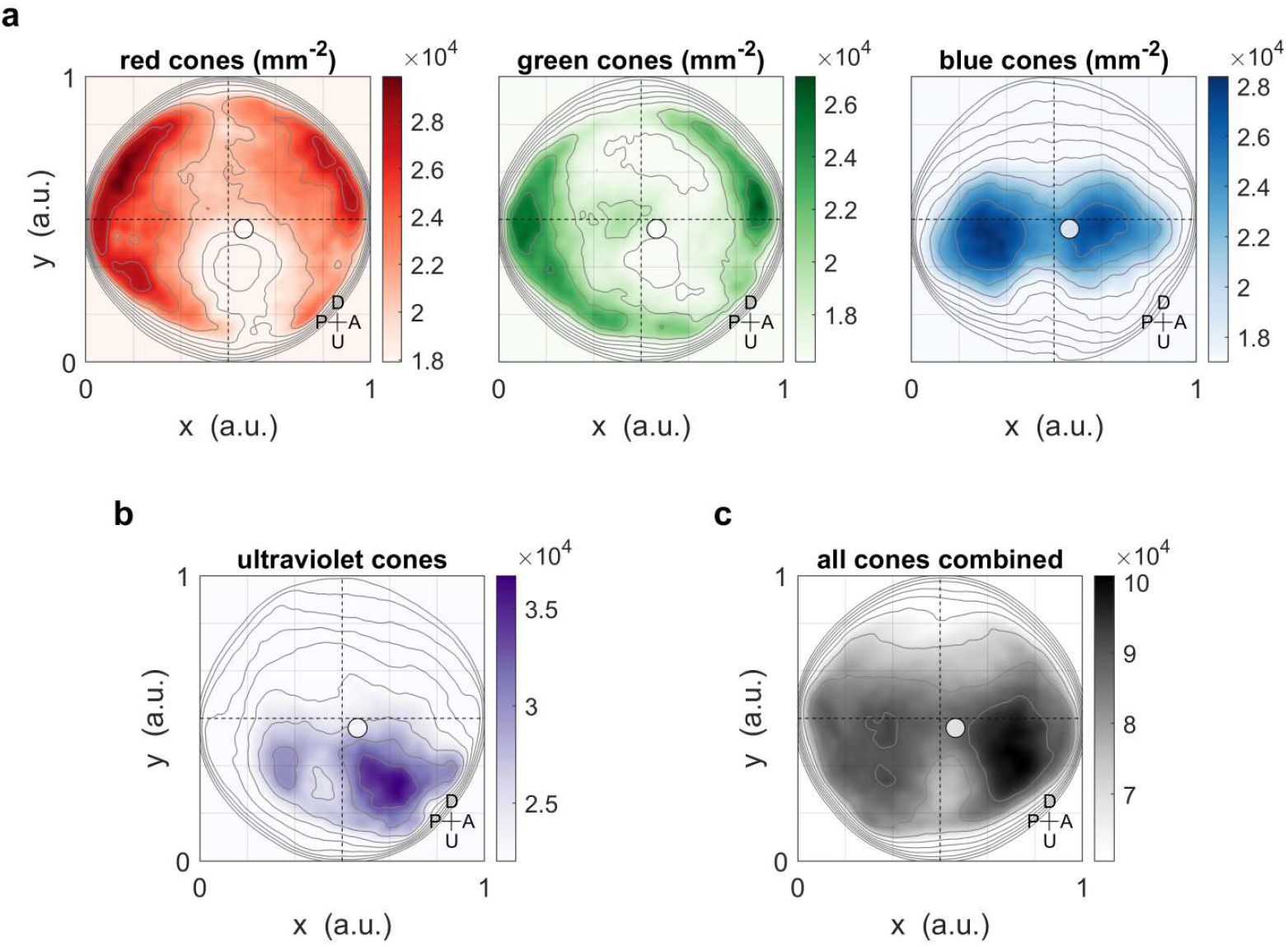
Maximum OKR gain compared to photoreceptor densities in retinal coordinates. Same as **Figure 6**, but showing positions across the retina in Cartesian coordinates, as originally published (Zimmermann et al. 2018 Curr Biol), instead of in geographic visual field coordinates. When plotting photoreceptor densities in Cartesian coordinates, the regions of highest densities appear to be located quite peripheral/eccentric. However, the plot of the densities in visual field coordinates **(Figure 6)** confirms the coincidence of high densities and high OKR gains. Solid circles indicate the location of maximum OKR gain inferred from experiments of type D in 5-7dpf larvae **(Figure 3)**, and corrected by the mean eye position over time.

### Supplementary Tables

**Supplementary File 4.** Arena cross-section. Elevation of LED tile centres, from top ribbon (ribbon number +5, near the north pole of the sphere) to bottom ribbon (ribbon number −5, near the South Pole). See **Figure 1–figure supplement 1a** for a graphical illustration. *As the left and right hemispheres of the arena are mirror-symmetric to one another, each ribbon contains the same number of tiles within each of the two hemispheres. **Because the structural scaffold is reinforced near the bottom of the sphere, ribbon −4 contains one fewer LED tile than ribbon +4. Other than that, the arrangement of LED tiles is almost mirror-symmetric from top to bottom as well.

| ribbon no. k | tiles per ribbon | elevation $\beta_k$ of tile centre (degrees) |
| --- | --- | --- |
| +5 | 2* x 5 | 62.4 |
| +4 | 2 x 9** | 49.9 |
| +3 | 2 x 11 | 37.6 |
| +2 | 2 x 13 | 25.0 |
| +1 | 2 x 14 | 12.5 |
| 0 (equator) | 2 x 15 | 0 |
| -1 | 2 x 14 | -12.5 |
| -2 | 2 x 13 | -25.0 |
| -3 | 2 x 11 | -37.6 |
| -4 | 2 x 8** | -49.9 |
| -5 | 2 x 5 | -62.4 |

**Supplementary File 5.** Absolute positions of fish and setup elements in different experiments. All positions and directions are given in environmental coordinates, i.e. the approximate cardinal directions (North etc.) of the laboratory, as well as Up and Down (away from or towards the core of the Earth). Most experiments, including control experiments, are of type 1 **(Figure 4a)**. The only exceptions are the position-dependence experiments with rotated arena (type 2, **Figure 4b**), control experiments with upside-down embedding (type 3, **Figure 4c**), and control experiments with inverted IR illumination (type 4, **Figure 4d**). Three LED tiles in different parts of the visual field are included as examples. Positions are given in lab-centred geographic coordinates as (a,e), where a is the azimuth, e the elevation, and 0° azimuth is arbitrarily chosen to point South.

| experiments | type 1 | type 2 | type 3 | type 4 |
| --- | --- | --- | --- | --- |
| **infrared illumination at** | Up<br>(0°, +90°) | Up<br>(0°, +90°) | Up<br>(0°, +90°) | Down<br>(0°, -90°) |
| **camera at** | Down<br>(0°, -90°) | Down<br>(0°, -90°) | Down<br>(0°, -90°) | Up<br>(0°, +90°) |
| **flat face of glass triangle facing** | Up<br>(0°, +90°) | Up<br>(0°, +90°) | Up<br>(0°, +90°) | Up<br>(0°, +90°) |
| **dorsum facing** | Up<br>(0°, +90°) | Up<br>(0°, +90°) | Down<br>(0°, -90°) | Up<br>(0°, +90°) |
| **rostrum facing** | South<br>(0°, 0°) | North<br>(180°, 0°) | South<br>(0°, 0°) | South<br>(0°, 0°) |
| **arena axis A** | South<br>(0°, 0°) | North<br>(180°, 0°) | South<br>(0°, 0°) | South<br>(0°, 0°) |
| **arena axis B** | West<br>(90°, 0°) | West<br>(90°, 0°) | West<br>(90°, 0°) | West<br>(90°, 0°) |
| **arena axis C** | Up<br>(0°, +90°) | Down<br>(0°, -90°) | Up<br>(0°, +90°) | Up<br>(0°, +90°) |
| **LED tile #1** | Up, West, South<br>(+35°, +53°) | Down, West, North<br>(+145°, -53°) | Up, West, South<br>(+35°, +53°) | Up, West, South<br>(+35°, +53°) |
| **LED tile #2** | East<br>(-90°, 0°) | East<br>(-90°, 0°) | East<br>(-90°, 0°) | East<br>(-90°, 0°) |
| **LED tile #3** | Down, East, South<br>(-16°, -25°) | Up, East, North<br>(+164°, +25°) | Down, East, South<br>(-16°, -25°) | Down, East, South<br>(-16°, -25°) |

**Supplementary File 6.** Stimulus parameters (control experiments). These stimuli consisted of a horizontally moving grating displayed on four flat, rectangular stimulus screens surrounding the larva. One pair of screens displayed stimuli visible to the left eye only, and the other pair displayed stimuli to the right eye only. Results shown in **Figure 3–figure supplement 5**.

| type | left eye stimulus | right eye stimulus | shown on panel |
| --- | --- | --- | --- |
| **C0** | moving pattern | moving pattern | **a-c** |
| **C1** | stationary pattern | moving pattern | **a-b** |
| **C2** | moving pattern | stationary pattern | **a-b** |
| **C3** | blank surface | moving pattern | **c-d, g-h** |
| **C4** | moving pattern | blank surface | **c-d, g-h** |
| **C5** | eye shielded | moving pattern | **e-f, g-h** |
| **C6** | moving pattern | eye shielded | **e-f, g-h** |

**Supplementary File 7.** Stimulus parameters (whole-field and hemispheres). These stimuli consisted of a horizontally moving grating, either covering the entire visual field or cropped to one of the 6 principal hemispheres (front, rear, upper, lower, left, right). The stimulus mask is determined by the azimuth *α* (degrees) and elevation *β* (degrees) of its centre, as well as its size, given by the angle *δ* (degrees) it spans. The moving grating is characterised by its spatial frequency SF (cycles/degree), temporal frequency TF (cycles/sec), peak velocity v (deg/sec), and oscillation period T (sec).

| type | $\alpha$ | $\beta$ | $\delta$ | SF | TF | $v$ | T |
| --- | --- | --- | --- | --- | --- | --- | --- |
| **H0** | 0 | 0 | 360 | 0.0611 | 0.7639 | 12.5 | 10 |
| **H1** | 0 | 0 | 180 | 0.0611 | 0.7639 | 12.5 | 10 |
| **H2** | 180 | 0 | 180 | 0.0611 | 0.7639 | 12.5 | 10 |
| **H3** | 0 | 90 | 180 | 0.0611 | 0.7639 | 12.5 | 10 |
| **H4** | 0 | -90 | 180 | 0.0611 | 0.7639 | 12.5 | 10 |
| **H5** | 90 | 0 | 180 | 0.0611 | 0.7639 | 12.5 | 10 |
| **H6** | -90 | 0 | 180 | 0.0611 | 0.7639 | 12.5 | 10 |

### Supplementary Videos

**Video 1.** (stimulusdistribution.avi) Repulsion-based algorithm to numerically distribute stimulus centres across a sphere surface **(Supplementary Code 2)**, with gradual convergence on a logarithmic timescale (blue to green). A subset of stimulus centres is held on the equator at zero elevation (orange).

**Video 2.** (locationstimulus.avi) Animation showcasing short samples of all disk stimuli used to study location dependence, as in **Figure 3a**.

**Video 3.** (frequencystimulus.avi) Animation showcasing short samples of all disk stimuli used to study frequency dependence, as in **Figure 5a**.

**Video 4.** (sizestimulus.avi) Animation showcasing short samples of all disk stimuli used to study size dependence, as in **Figure 5b**.

